# Eco-evolutionary significance of ‘loners’

**DOI:** 10.1101/508507

**Authors:** Fernando W. Rossine, Ricardo Martinez-Garcia, Allyson E. Sgro, Thomas Gregor, Corina E. Tarnita

## Abstract

Loners, individuals out-of-sync with a coordinated majority, occur frequently in nature. Are loners incidental byproducts of large-scale synchronization attempts or are they part of a mosaic of life-history strategies? Here, we provide the first empirical evidence of naturally occurring heritable variation in loner behavior, using the social amoeba *Dictyostelium discoideum*. Moreover, we show that *Dictyostelium* loners—cells that do not join the multicellular life-stage— result from a dynamic population-partitioning process. Underlying this partitioning, we find evidence that each cell makes a stochastic, signal-based decision resulting in an imperfectly synchronized multicellular development affected by both abiotic (environmental porosity) and biotic (strain-specific signaling) factors. Finally, we predict that when strains differing in their partitioning behavior co-occur, cross-signaling impacts slime-mold diversity across spatio-temporal scales. Loners are therefore critical to understanding collective and social behaviors, multicellular development, and ecological dynamics in *D. discoideum*. More broadly, across taxa, imperfect synchronization might be adaptive by enabling diversification of life-history strategies.

## Introduction

Collective behaviors, in which a large number of individuals exhibit some degree of behavioral synchronization, are frequent across the tree of life and across spatio-temporal scales: from microbial aggregates to the great wildebeest migration, from locust swarming to synchronized bamboo flowering, from fish schooling to mechanical adaptation in honeybee clusters (Couzin & Krause, 2003; Gregor, Fujimoto, Masaki, & Sawai, 2010; Hopcraft et al., 2015; Janzen, 1976; Kaiser & Crosby, 1983; Katz, Tunstrom, Ioannou, Huepe, & Couzin, 2011; Peleg, Peters, Salcedo, & Mahadevan, 2018; Simpson, McCaffery, & Hägele, 1999). Intriguingly, however, such synchronization is sometimes imperfect and ‘out-of-sync’ individuals (henceforth loners), have been reported in several of these systems. For instance, in locusts, population crowding prompts a transition from a solitary phase, in which individuals repel each other, to a gregarious phase, in which they attract each other. Experiments show, however, that not all individuals undergo this transition, even if exposed to long periods of crowding (Simpson et al., 1999). In wildebeest, hundreds of thousands of individuals coordinate with each other and organize herding migrations, but resident populations, that fail to migrate, also exist (Hopcraft et al., 2015). Similarly, wildebeest calving times are also highly coordinated, but some fraction of the calves are born outside the calving period (Hopcraft et al., 2015). In bamboo, individuals predominantly flower in synchronized masts, but sporadic out-of-sync events have also been recorded (Janzen, 1976).

The roots of imperfect synchronization will undoubtedly differ across systems. Nevertheless, the occurrence of imperfect synchronization across such different systems and scales raises fundamental questions about its causes and consequences. Are loners mistakes—merely inevitable byproducts of large scale synchronization attempts—or are they a variable trait that selection can shape with potential ecological consequences? Theoretical investigations of such loner behaviors have been sparse, but the handful of existing studies have suggested that, at least in some systems, they could be a means of spatio-temporal niche-partitioning (Dubravcic, van Baalen, & Nizak, 2014) that promotes diversity (Martínez-García & Tarnita, 2017; Tarnita, Washburne, Martínez-García, Sgro, & Levin, 2015). However, despite this theoretically established potential, variability and heritability of loner behaviors have not been characterized in natural populations. Thus, there exists no empirical evidence, in any system, that loners are anything more than chance stragglers, lacking an avenue for selection to act on them.

The cellular slime mold *Dictyostelium discoideum* is an ideal system in which to experimentally characterize loner behaviors. Its life cycle comprises a unicellular feeding stage and a starvation-induced multicellular stage—the result of a developmental process involving coordinated cell aggregation, which culminates in the production of starvation-resistant spores (Bonner, 2009). There has been extensive progress in understanding this multicellular stage (Bonner, 2009; Gregor et al., 2010; Strassmann & Queller, 2011), but less attention has been paid to the potential role of asocial aspects—such as the non-aggregating solitary loner-cells—in development. Loners die under sustained starvation, but they persist temporarily (Dubravcic, 2013); if food is replenished, they eat and divide, and their progeny subsequently recapitulate the multicellular development (Tarnita et al., 2015). Finally, it has been shown that knockout strains can have different loner behaviors (Dubravcic et al., 2014). Altogether, these observations suggest that loners could indeed be part of a life-history strategy in *D. discoideum*. However, to fully establish this, one needs to show (a) that there are evolutionary paths such that loner behavior can be tuned while populations retain viability in their natural environments and (b) that there are fitness differences between strains with different loner behaviors in natural environments. Here we tackle (a) by inspecting loner behavior in naturally occurring strains; in the process, we uncover the cellular decision-making rules underlying the aggregator-loner partitioning and explore their potential ecological consequences.

## Results and Discussion

### The aggregator-loner partitioning is heritable and context-dependent

To determine whether the loner behavior is heritable—and, thus, whether there is the potential for natural selection to act on it—we developed an experimental protocol to identify individual loner cells (Fig. 1a,b), permitting us to characterize their spatial distribution (Fig. 1c, Fig. S1) and quantify their density (Fig. 1d,e, Fig. S2). Importantly, we worked with three natural isolates (i.e. strains or genetic variants) that were collected from the same location to ensure that observed behaviors of individual strains were not an artifact of lab rearing and that observed behavioral differences among strains reflected naturally co-occurring strategies. When homogeneously plated cells of a given strain were left to starve and undergo multicellular development, loner cells were found throughout aggregation territories, with a higher density at territory borders and at experimental boundary conditions than in the immediate surroundings of aggregation centers (Fig 1c, Fig. S1). In repeated experiments under controlled conditions, loner densities of a given strain fell consistently within a conserved distribution (framed portion of Fig. 1e); moreover, the loner distributions of some strains were significantly different in their mean and variance (compare strains NC28.1 and NC85.2 in Fig. 1d, and framed portion of Fig. 1e). These findings demonstrate that the aggregator-loner partitioning behavior is heritable and thus has the potential to be shaped by selection.

**Figure 1.**
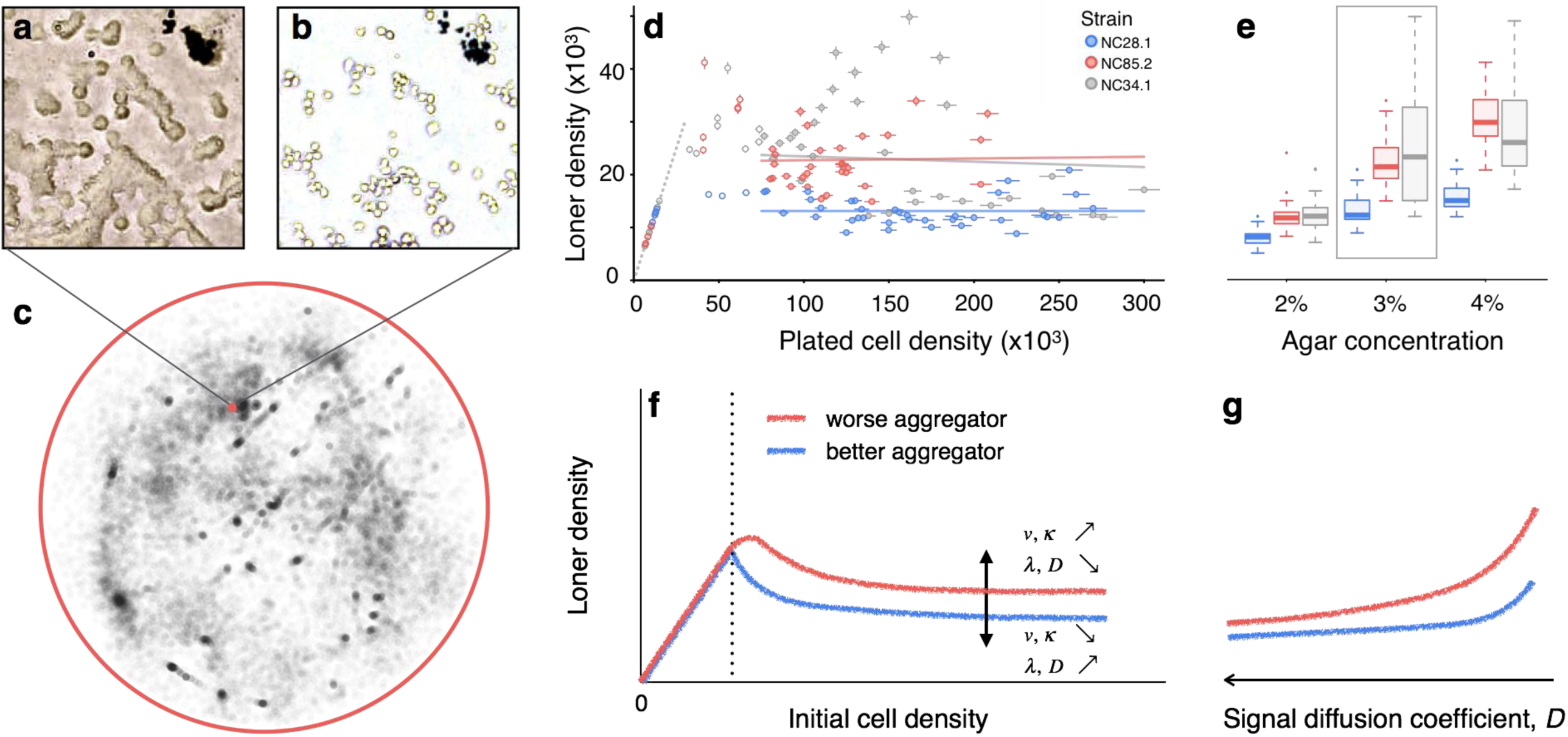
Loners are a heritable component of *D. discoideum* fitness. **a, b,** After aggregation, loner cells are hard to individualize (**a**), but become clearly distinguishable after processing (**b**). **c,** Map of the position of each loner cell in an experiment (NC85.2 developing in 3% agar). Red square marks region shown in (**a,b**). **d,** Loner densities of three strains as a function of initial plated density in 3% agar. Error bars, independent estimates of counting error (see Methods). Lines correspond to linear regressions using only high-initial-density data points (filled circles, >75.000 cells/cm^2^). **e,** Loner densities from experiments with high initial cell density as a function of substrate agar concentration (y-axis same as in **d**). Boxes, inter-quartile ranges; horizontal lines, medians; whiskers, 1.5× inter-quartile range from the median; points, outliers. Strain NC28.1 always left fewer loners (*t*-test, p<0.001). Values inside frame correspond to data used in (**d**). **f, g** Schematic of model results showing loner densities as a function of initial cell density (**f**) and signal diffusion coefficient (**g**). The y-axis in (**g**) is the same as in (**f**).

To characterize the underlying developmental process, it must first be determined whether a cell’s decision to commit to aggregation or remain a loner is context-independent (Dubravcic et al., 2014; Tarnita et al., 2015) (the result of a stochastic switch) or whether it depends on external factors. If it is context-independent, then loner density should increase linearly with the density of initially plated cells, i.e., the heritable quantity would be the fraction of loners, as previously posited (Dubravcic et al., 2014; Tarnita et al., 2015). Instead, we found a non-linear dependence: at low initial densities, cells were too sparse for aggregation to occur and all cells remained loners; above a threshold, aggregation occurred with increasing efficiency and loner densities decreased; surprisingly, at high initial cell densities, loner densities plateaued (Fig. 1d, Fig. S2c-k). Thus, past a range of initial densities, it is the *number* (or density), and not the *fraction,* of loners that is heritable, suggesting an underlying cell decision-making process that is fundamentally different from a stochastic switch. Furthermore, when we varied the porosity of the agar substrate—a proxy for an important environmental characteristic for this soil-dwelling amoeba—less porous (more concentrated) agar yielded higher loner densities (Fig. 1e), and differentially affected *D. discoideum* strains by enhancing the difference between strains that leave fewer loners (‘better aggregators’) and those that leave more loners (‘worse aggregators’). Interestingly, the agar porosity affects not only the mean number of loners, but also the variance (Fig. 1e). Altogether, these findings demonstrate that the heritable aggregator-loner partitioning is context-dependent— the result of a density-dependent decision-making process (Balázsi, Van Oudenaarden, & Collins, 2011) that interacts with the abiotic environment.

### The aggregator-loner partitioning is the result of an abiotically-modulated quorum-based stochastic process

To identify the properties of this decision-making process, we constructed a spatially explicit individual-based model (Fig. S3; see Methods) starting from a limited set of assumptions: that the strain-specific partitioning behavior results from imperfect synchronization in the developmental program, and that this asynchrony stems from a stochastic response to cell-population densities (i.e., quorum sensing). Consistent with our experimental design, we started with a population of cells immediately after food exhaustion and assumed them to be in a pre-aggregating state, *P*. *P*-cells emit extracellular signaling molecules at a strain-specific rate γ; signal diffuses with diffusion coefficient *D* and serves a quorum-sensing purpose (Gregor et al., 2010; Loomis, 2014) that regulates the stochastic transition to the aggregating state, *A*: when the signal perceived by a cell exceeds the strain-specific sensitivity threshold *θ* (i.e. the quorum is met), that cell has a strain-specific probability λ per unit time of becoming an aggregating *A*-cell. *A*-cells continue to emit signal and move towards the aggregation center with constant, strain-specific velocity 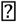. At the center, cells become multicellular (*M*-state) and stop emitting signal. Importantly, we do not impose restrictions on the nature of the signaling molecule or on the sensing mechanism, and the model employs this molecule broadly to fulfill both a traditional (Gomer, Yuen, & Firtel, 1991; Loomis, 2014) and a dynamical quorum sensing role (Gregor et al., 2010).

Because our model is focused on population partitioning, it deliberately simplifies certain dynamics (Dallon & Othmer, 1997; Gregor et al., 2010; Keller & Segel, 1970; Kessler & Levine, 1993; Palsson & Othmer, 2000; Sawai, Thomason, & Cox, 2005) and it bundles many possible intermediate states that make up the *P*-to-*A* transition: for example, before aggregating, cells must sequentially starve, become excitable by cAMP (Clarke & Gomer, 1995; Jain, Yuen, Taphouse, & Gomer, 1992), and finally chemotax (Gregor et al., 2010). Moreover, for computational convenience, our model simplifies various sources of stochasticity that arise during the developmental process, such as the existence of multiple aggregation territories and the variable boundary shapes between them. This prevents us from studying the sources and dynamics of the variance in loner density, an important direction for future work. Nevertheless, this reduced model is powerful enough to qualitatively recapitulate all other properties of the observed population partitioning (Fig. 1f; Fig. S4a-e). Importantly, the model recovers the plateau in the loner counts as a function of initially-plated cell density (Fig. 1f; Fig. S4a-d; Box 1 and SI Appendix) and it provides an intuition for the identity of the loners (Fig. 2): they are *P*-cells that did not make the probabilistic transition to the *A*-state when they had a quorum and that are left without a quorum when enough of their neighbors underwent the *P*-to-*A* transition and moved towards the aggregation center.

**Figure 2.**
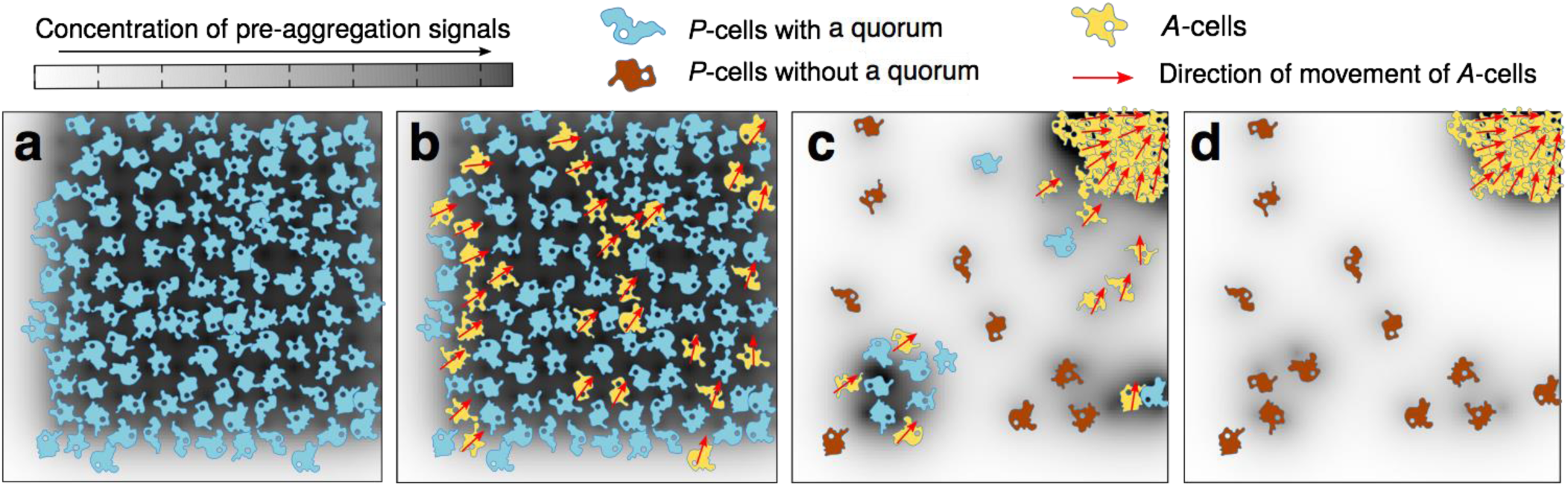
Developmental model schematic. **a,** At high initial densities, all *P*-cells have a quorum to initiate aggregation. **b,** With a strain-specific probability, some *P*-cells transition into *A*-cells. **c,** As *A*-cells aggregate, some of the *P-*cells that did not transition into *A-*cells are left without a quorum. These are the loners. **d,** At the end of development, *P-*cells far from the aggregate location are more likely to have been left without a quorum and to stay as loners.

Total loner density then depends on how quickly *P*-cells switch to the *A*-state relative to how quickly they are left without a quorum (λ*/v*), and on how easy it is to maintain a quorum (see Fig. S4a-d, Box 1 and SI Appendix for analytical results). Thus, the larger the *P*-to-*A* transition rate λ, the fewer loner cells are left behind since *P*-cells sensing a quorum switch faster to the *A*-state; conversely, the larger the aggregation speed *v*, the more loners are left behind since *A*-cells move away faster and leave their *P*-cell neighbors without a quorum (Fig. S4c). Consequently, the farther a cell is from the aggregation center, the sooner it is left without a quorum and the more likely it is to become a loner (Fig. S4e), which explains our experimentally observed spatial distribution (Fig. 1c).

Achieving and maintaining a quorum depends on the ratio between the sensitivity threshold and the signaling rate, *κ* = *θ/γ*, and on the signal diffusivity, *D*. Higher *κ* leads to more loners because more neighbors are required for a quorum (Fig. S4d). Similarly, lower diffusivities result in higher loner densities (Fig. S4f,g) because the signal remains highly concentrated around the emitters, and cells need to be more densely packed to maintain a quorum (Fig. S4h,i). Moreover, decreasing the diffusivity differentially affects worse and better aggregators (Fig. 1g, Fig. S4f,g), because diffusivity and signal spreading are nonlinearly related (see Methods). These results mirror the experimentally determined dependence of loner densities on agar concentration, suggesting signal diffusivity as a potential mediator of this dependency. Because loner densities responded to agar-concentration changes in a range that should not impede the diffusion of cAMP (Johnson, Berk, Jain, & Deen, 1996; Pluen, Netti, Jain, & Berk, 1999), these results further suggest that at least one of the molecules involved in the quorum-dependent transition should be large—for example, conditioned medium factor (CMF)(Gomer et al., 1991), prestarvation factor (PSF) counting factor (Kolbinger et al., 2005), counting factor (Brock & Gomer, 1999) or phosphodiesterase (Bodenschatz, Bae, & Prabhakara, 2017).

Notably, PSF and CMF are secreted during the growth phase and early starvation. This led us to investigate the potential role that these earlier signaling stages could play in regulating loner behavior. To test this hypothesis, we let cells grow in bacterial suspension until resources are depleted, and only subsequently plated them in agar gels. Thus, the initial responses to resource depletion occur in a well-mixed environment, and any signaling molecules secreted in this phase should synchronously reach all cells. If early signaling is responsible both for the loner differences between strains and for the effects of agar concentration on loners, we predicted that the well-mixed environment should produce the same effects as increasing diffusion in our model (Fig 1g). First, the increased signaling synchrony should decrease the loner number of any given strain; second, strains that had more synchronous signaling to begin with (i.e. better aggregators) should be less affected by this treatment than strains that started out with less synchronous signaling (i.e. worse aggregators), which would cause the differences between strains to decrease. Consistent with these predictions, we found that the loner counts decreased dramatically for the worse aggregator, leading to a reduction in the difference between strains (Fig. 3). This supports our hypothesis that vegetative or early starvation signaling—and not the later, cAMP relay signaling and synchronization, as previously inferred using knockouts (Dubravcic et al., 2014)—could be a critical stage at which loner behavior is regulated and the natural variation that we observed is produced.

**Figure 3.**
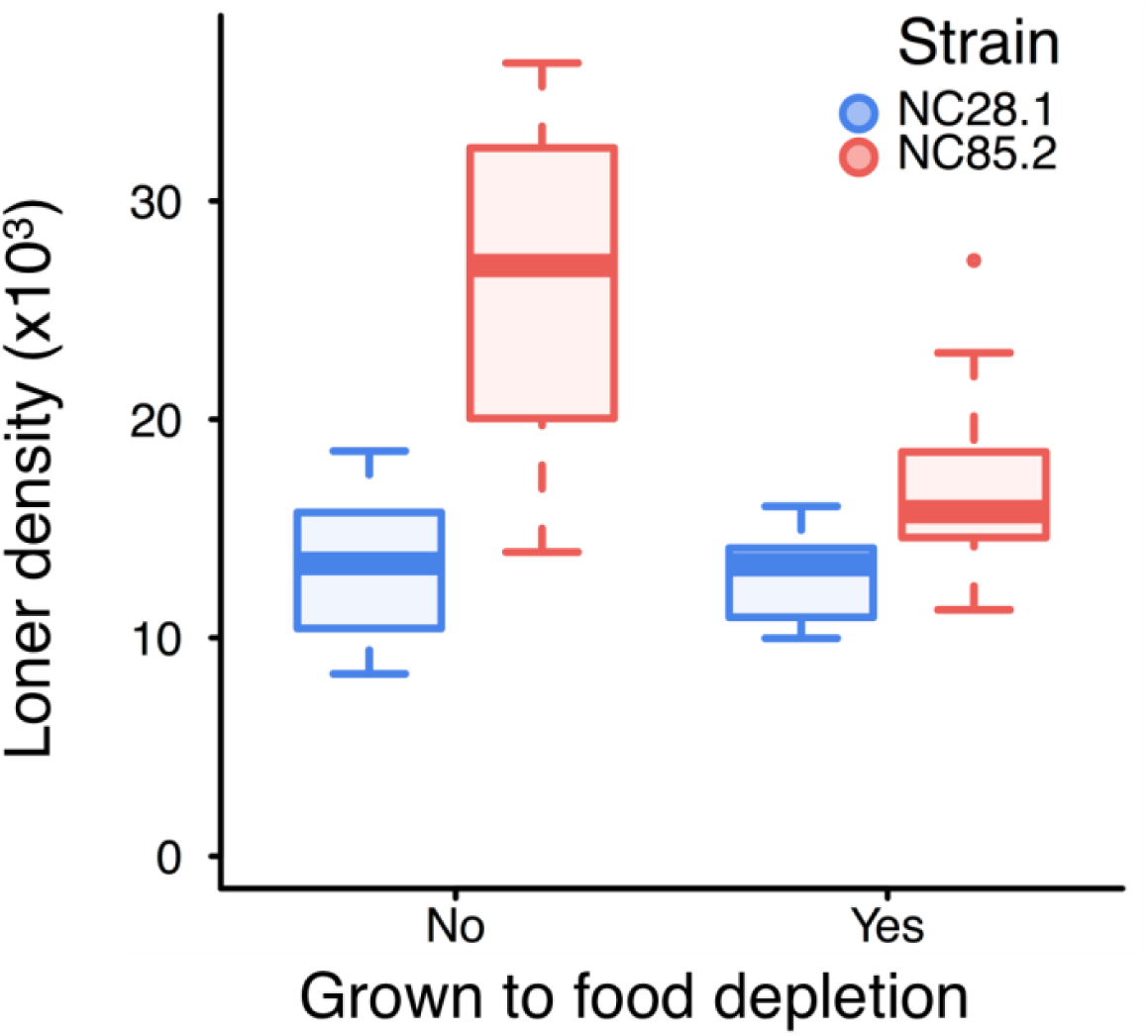
Well-mixed signaling changes loner behavior. Allowing cells to exhaust their resources in suspension does not significantly change the behavior of the better aggregator (strain NC28.1) (p = 0.7), but it reduces the mean and variance of the number of loners of the worse aggregator (strain NC85.2) (p = 0.0006). Boxes, inter-quartile ranges; horizontal lines, medians; whiskers, 1.5× inter-quartile range from the median; points, outliers.

### The aggregator-loner partitioning depends on the identity of neighboring cells

Collectively, the results above show that the population partitioning stems from interactions between genotype and environment and suggest that cell signaling mediates these interactions. This raises the possibility that a strain’s partitioning could also be influenced by the presence of other strains via cross-signalling. If a cell’s commitment to aggregation were independent of the identity of co-occurring strains, a mix of strains would leave behind a total mixed loner density that is the linear combination of the two strains’ loner densities (see Methods). Our model, however, predicts developmental interactions between co-occurring strains that produce a diversity of departures from linearity (Fig. 4a; Fig. S5).

**Figure 4.**
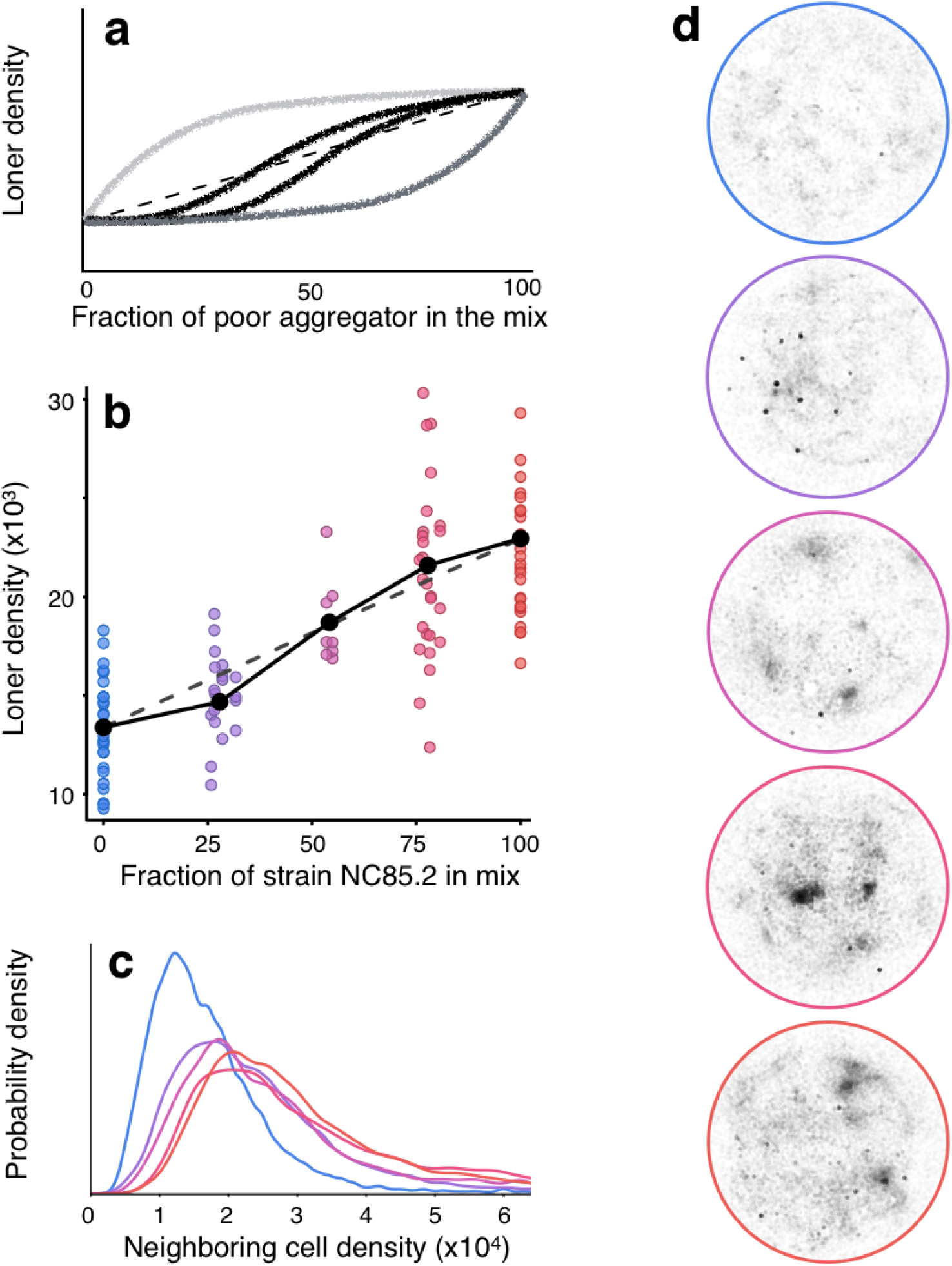
Co-occurring strains interact during development. **a,** Schematic of the range of theoretically predicted loner densities at different mix proportions of two co-occurring strains (thick curves). Dashed line, expected loner densities if cells commit to aggregation independent of the identity of their neighbors. **b,** Experimentally observed loner densities for different mix proportions of a better (NC28.1, blue) and a worse (NC85.2, red) aggregating strain in 3% agar. Black points, mean loner densities for each of the five proportions. Dashed line, same as in (**a**). Solid lines, piecewise linear regressions, which deviate significantly from the linear fit (p=0.019; see Methods). **c,** Experimentally observed spatial patterns of each mix proportion are characterized by the local cell density around each cell (see Methods). Narrower distributions (NC28.1, blue curve) correspond to more homogeneously distributed loners. Broader distributions (NC85.2, red curve) correspond to more clumped loners. **d**, Experimentally obtained loner position maps are shown for each of the mixed proportions. Colors in (**c,d**) correspond to colors in (**b**).

When we plated well-mixed cells of the strains NC28.1 (better aggregator) and NC85.2 (worse aggregator) at different frequencies and left them to co-develop under starvation conditions we found agreement with this theoretical prediction. The total loner density of the mixed strains deviated significantly from the linear combination, mapping out a sigmoidal curve (Fig. 4b, Fig. S6), which was one of three possible theoretical outcomes. Thus, when the better aggregator was more abundant in the mix (25%:75%) there were fewer total loners than predicted by the linear combination; conversely, when the worse aggregator was more abundant (75%:25%) there were more total loners. That strains influence each other’s partitioning is consistent with existing results using knockouts (Dubravcic et al., 2014) and it is particularly interesting in light of prior work showing that, during aggregation, *D. discoideum* cells do not perfectly discriminate against non-kin and genetically heterogeneous multicellular aggregates occur naturally (Strassmann, Zhu, & Queller, 2000), allowing for potential interactions between strains that can alter each other’s life-history investments (Buttery, Rozen, Wolf, & Thompson, 2009; Strassmann & Queller, 2011). Whether or not such interactions occur within the aggregate (Martínez-García & Tarnita, 2016; Tarnita et al., 2015; J. B. B. Wolf et al., 2015), our results reveal that they do occur earlier in the developmental process.

Such developmental interactions that alter life history investments could severely impact strain fitness, alter *D. discoideum* diversity, and threaten the persistence of the social behavior (Strassmann & Queller, 2011). It is therefore crucial to understand their consequences for individual strains. In our case, the theoretical model produced two possible outcomes of co-development (see SI Appendix): the two co-occurring strains can become either (i) more similar (Fig. S7a,b) or (ii) more different (Fig. S7c,d) in their partitioning behavior. In particular, when the theoretical density of the mixed-strain loners had a sigmoidal shape similar to that derived experimentally, the loners of the better aggregator quickly went to zero as the frequency of the worse aggregator in the mix increased (Fig. S7c,d). Thus, the better aggregator became even better in the presence of the worse aggregator, enhancing the difference between the two strains (case ii). This occurred because the more sluggish loners of the worse aggregator maintained quorum long enough for the better aggregator to aggregate perfectly. Experimentally, the spatial distribution of the mixed-strain loners provides insight into their potential composition (Fig. 4c,d): as soon as the worse aggregator is part of the mix (even at the lowest frequency), the spatial distribution of the mixed loners is almost identical to that of the worse aggregator—and strikingly different from that of the better aggregator—suggesting that, as predicted, the mixed-strain loners predominantly comprise the worse aggregator. Importantly, developmental interactions between co-occurring strains do indeed have consequences for the life history investments of individual strains.

### Cross-signaling may foster slime mold diversity across spatio-temporal scales

The two possible outcomes of co-development predicted by our population-partitioning model are reminiscent of two classical evolutionary routes to diversity maintenance—quasi-neutrality (case i; Fig. 5a) and character displacement (case ii; Fig. 5b)—and are therefore likely to have consequences for slime-mold diversity. To investigate these biodiversity consequences of strains mixing and co-developing—instead of perfectly segregating and avoiding co-development—we incorporated our population-partitioning model into an existing model of competition for resources over multiple successive growth-starvation (Martínez-García & Tarnita, 2016, 2017; Tarnita et al., 2015) (Fig. S8; see Methods). Although empirically we did not investigate fitness differences between strains, here we assume that such differences exist and depend on the environmental conditions as in prior work (Martínez-García & Tarnita, 2016, 2017; Tarnita et al., 2015). The environment is characterized by the mean time between nutrient replenishment events. We considered both deterministic environments (all replenishment times of equal size) and stochastic environments (exponentially distributed nutrient replenishment times). For each, we explored a range of mean nutrient replenishment times. Regardless of whether co-development occurs, within any environment, there was competitive exclusion: consistent with previous work, we found that strains that leave behind more (fewer) loners are more competitive in environments with shorter (longer) mean replenishment times (Tarnita et al., 2015). In deterministic environments the identity of the winner was also deterministic and not altered by co-development (inset of Fig. 5c,d); however, co-development did alter the time-to-extinction of the loser (inset of Fig. 5e,f). On the contrary, in stochastic environments, for every pair of competing strains, there is a range of environments where the identity of the winner is uncertain and that range is drastically altered by co-development (Fig. 5c,d). As in deterministic environments, co-development also influenced the time-to-extinction of the loser (Fig. 5e,f).

**Figure 5.**
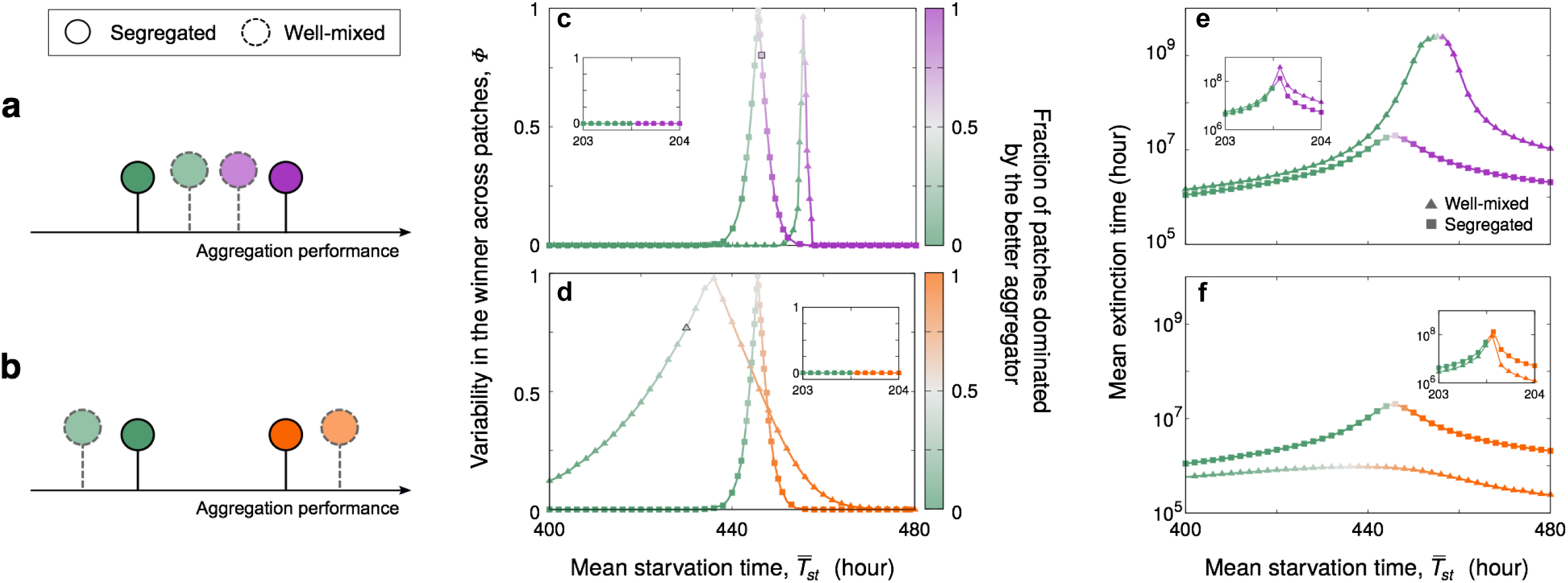
Ecological implications of developmental interactions. **a, b,** Co-development makes strains (different colors) more similar (**a**, case i) or more different (**b**, case ii) in their partitioning behavior. **c-f**, Effect of well-mixed versus segregated development on (**c, d**) the identity of the winner and (**e, f**) extinction times. Theoretical dynamics in the two environments whose symbols have a black boundary are shown in Fig. 6. Panels **c, e** and **d, f** are predictions for the pairs of strains in Figures S7a and S7c, respectively. Green strain is the same across all panels. Main panels, stochastic environments; insets, deterministic environments (see Methods). Color gradients indicate the change in the fraction of patches dominated by each of the mixed strains. Parameterization as in Fig. S7a,c

To untangle the effects of co-development on diversity at different spatial scales, we discretized the environment into small-scale patches with identical replenishment conditions. Competition between pairs of strains occurred within each patch, and there was no dispersal between patches. Importantly, this setup allows us to investigate the effects of co-development on alpha (intra-patch) and beta (inter-patch) diversity, but it does not introduce any intrinsic spatial heterogeneity. As expected, within each patch, we found competitive exclusion. However, at the level of the environment, if the replenishment conditions are within the range where the identity of the winner is uncertain, there can be coexistence. Each of our predicted modes of co-development imparted a distinct biodiversity signature. Specifically, in case (i), the converging behaviors of the two strains led to much longer times to extinction, resulting in higher transient alpha diversity compared to the segregated model. However, this mode of co-development also narrowed the environmental range in which competition lead to non-deterministic exclusion, resulting in lower stationary beta diversity compared to the segregated model (Fig. 6a). Case (ii) yielded the opposite outcome for both alpha and beta diversity (Fig. 6b).

**Figure 6.**
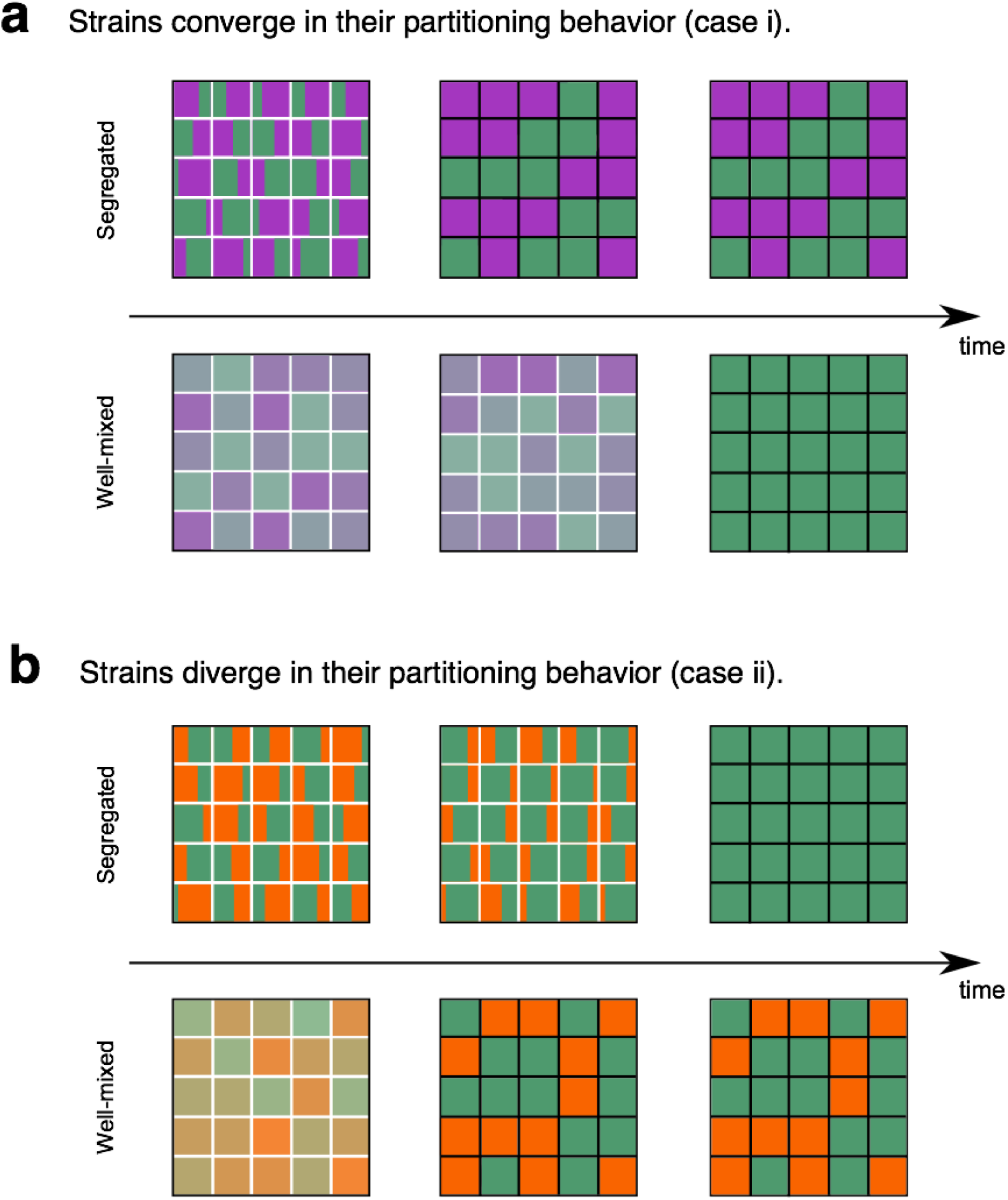
Model results of the effects of developmental interactions on transient alpha-diversity and stationary beta-diversity. **a,b,** White interior boundaries symbolize a transient patch-level dynamic; black interior boundaries symbolize a stationary patch-level dynamic. a, Strains converge in their partitioning behavior due to developmental interactions (case i). In a stochastic environment with mean starvation time 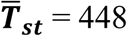 hour (symbol with black boundary in Fig. 4e), co-development induced quasi-neutrality increases transient coexistence (and transient alpha-diversity). However, it also eliminates the variability in the winner and thus co-development induced quasi-neutrality eliminates stationary beta-diversity. b, Strains diverge in their partitioning behavior due to developmental interactions (case ii). In a stochastic environment with mean starvation time 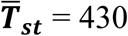 hour (symbol with black boundary in Fig. 4e), co-development induced character displacement has the opposite effect from (a): the mean extinction time decreases (decreased transient alpha-diversity) but the variability in the winner across patches increases compared to segregated development (increased stationary beta-diversity).

## Conclusion

To conclude, here we showed natural variation and heritability in the aggregator-loner partitioning behavior of naturally co-occurring strains of *D. discoideum*. Strikingly, the seemingly asocial loners are not a separate, independently-determined subset of cells, but rather they arise dynamically from the collective process. Coupling experiments and theory, we revealed that the aggregator-loner partitioning behavior is governed by a stochastic cell-level decision-making process mediated by cell signaling and modulated by both the abiotic and the biotic context. These investigations of collective behavior revealed previously unknown stochastic aspects of *D. discoideum* development. Finally, we used a theoretical approach to explore the ecological consequences of these findings and showed that the co-development of different strains impacts diversity across multiple scales. These results, arising solely from interactions between ecology and development, recapitulate the biodiversity outcomes of classical eco-evolutionary interactions. Overall, our results highlight the necessity of an integrated approach to collective behaviors, including multicellularity: studying asocial life-history strategies revealed insights into collective behavior and development, and studying development revealed insights into ecological dynamics and parallels with well-studied eco-evolutionary processes.

Furthermore, our findings have potentially fundamental implications for the evolution of aggregative multicellularity and social behaviors. Many factors could undermine the integrity of social complexes, such as free-riding, whereby individuals reap the benefits of group living without paying the costs. In slime molds, free-riders—strains that never contribute to stalk formation in mixes—have been found both in the wild (Buss, 1982) and in the lab (Kuzdzal-Fick, Fox, Strassmann, & Queller, 2011). If under selection, the heritable loners, invulnerable to the threats to the multicellular stage but capable of re-achieving multicellularity via their offspring, could constitute insurance against such threats and could therefore be critical to the evolution and persistence of aggregative multicellularity. This hypothesis is consistent with the handful of theoretical studies on cooperative behaviors that have considered social loners (Garcia, Doulcier, & De Monte, 2015; Hauert, De Monte, Hofbauer, & Sigmund, 2002).

Finally, beyond multicellularity and sociality, our results have potential implications for the broadly analogous loner behaviors identified across a variety of systems in which some form of coordination or synchronization is observed, from insects (Simpson et al., 1999) to vertebrates (Couzin & Krause, 2003; Hopcraft et al., 2015) to plants (Janzen, 1976). Our findings represent the first demonstration that loner behaviors can indeed exhibit natural variation and heritability— and that this can have significant ecological consequences—and, as such, they motivate a broader investigation into loner behaviors in other systems. While the mechanisms underlying the existence of loners are likely different across systems, the widespread existence of loners and the possibility that they could in fact be shaped by selection suggest an interesting conjecture: that, in general, imperfect synchronization may enable evolution to shape population-partitioning strategies in ways that could be instrumental for behavioral diversity and for the persistence of the collective stage, and thus for system-level robustness.

#### Box 1

In the spatially implicit limit (*D* ⟶ *∞*, *v* finite) and in the limit of large initial population size (*N*_0_ ⟶ *∞*), the density of loners *ρ*_*L*_ can be calculated analytically, which reveals the interactions among model parameters (see SI Appendix). We find

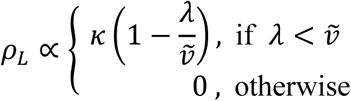

where *κ* = *θ/γ* is the ratio between the sensitivity threshold and the signaling rate, and 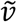 is the velocity rescaled by the mean distance traveled by cells before joining the aggregate. This reveals a phase separation determined by the relative speeds of the two transitions, *P*-to-*A* and *A*-to-*M* (Fig. S4b): a lack of synchronization resulting in loners occurs only if the *P*-to-*A* transition is slower than the *A*-to-*M* transition.

## Acknowledgments

We thank J. Bonner, E. Cox, D. Morris, D. Liberles, R. Pringle, M. Skoge, and K. Vetsigian for useful discussions. This work was supported by the Gordon and Betty Moore Foundation through grant GBMF2550.06 (RMG), NIH R01 GM098407 (TG), NRSA F32 GM103062 (AES), and the Alfred P Sloan Foundation (CET). AES was partially supported by a Burroughs Wellcome Fund Career Award at the Scientific Interface.

## Author contributions

All authors contributed to the conception of the study. FWR, AES and TG designed the experiments with input from RMG and CET. FWR performed the experiments with initial guidance from AES. FWR analyzed the experiments. RMG developed the theoretical models with input from FWR and CET. RMG performed and analyzed the simulations. RMG and CET produced analytical calculations. CET drafted the paper, and all authors provided comments.

## Data and materials availability

The data sets generated and analysed during the current study are available in the Dryad repository at http://dx.doi.org/XXX. The computer code is available from the corresponding authors upon request.

## Materials and Methods

### Experiments. *D. discoideum* growth and plating

NC28.1, NC34.1, and NC85.2 —three clonal lineages of *D. discoideum* originally isolated from Little Butt’s Gap, North Carolina (Francis & Eisenberg, 1993) –were obtained from dictyBase (Fey, Dodson, Basu, & Chisholm, 2013) and grown *on Klebsiella aerogenes* lawns prepared on SM agar plates (Fey, Kowal, Gaudet, Pilcher, & Chisholm, 2007). After the *D. discoideum* cells aggregated and formed fruiting bodies, spores were harvested and used to inoculate 3mL of a *K. aerogenes* suspension in SorMC buffer (OD600 of 8). The suspension was kept in a shaker for 24 hours and then used to inoculate 12mL of the *K. aerogenes* suspension. During growth*, D. discoideum* cell densities were kept below 3 x 10^6^ cells/mL. After 24 more hours, in preparation for the synchronous starvation of the cells, the suspension was cooled to 4°C for five minutes. The suspension was then centrifuged at 700 g for three minutes at 4°C and the remaining pellet was resuspended in 10mL of SorMC buffer. The spinning and resuspension cycles were repeated three times to wash away any remaining *K. aerogenes* cells (Fey et al., 2013). For the final resuspension the cell concentration was 10^7^ cells/mL.

From each stock suspension, a dilution series in SorMC buffer (80%,70%,…,20%) was obtained. In addition, a 5% dilution was prepared from each stock suspension. The 5% dilutions were below the critical aggregation density, and they were used to estimate the total amount of cells in the other samples coming from the same stock suspension. Cells were platen on non-nutrient agar gels (2%, 3% and 4% concentrations) cast in 1.5mm acrylamide gel casts (Bio-Rad). Each of the diluted and undiluted cell suspensions was applied to the agar substrates as a 10μL droplet.

The samples were then left to develop in a moist dark chamber at 21°C until the streaming phase of aggregation was over and most of the aggregates were already at the slug stage (∼14 hours). Development was then halted by lowering the chamber temperature to 4°C. Even though the 5% diluted suspension samples never aggregated, they were also left in the chamber for the full length of the experiment. This circumvents the problem of residual divisions post resource removal. The diluted and non-aggregated samples were then used to estimate the total number of cells in the undiluted aggregated samples.

In order to check for the consistency of the dilution procedure, 5%, 10% and 15% dilutions of NC34.1 strain stock solutions were also prepared and plated on 3% agar substrates. A linear Gaussian model with intercept 0 was fit to the cell counts of different dilutions of the same suspension. The standard deviation of this model was taken to be the error intrinsic to the dilution process (Figure S2a).

For mixed strain experiments, strains NC28.1 and NC85.2 were used. They were grown, washed and resuspended separately. Then, without diluting the suspensions, different mixes were made (25%,50% and 75% of the initial strain NC28.1 suspension). A 5% dilution was made for each of the two pure strain suspensions. Each of the mixes, pure strain suspensions, and dilutions were plated as four replicates for each experiment. These 5% dilutions were used to estimate the proportions of cells of each strain in each of the experiments. 3% agar substrates were used.

For the resource depletion experiments, cells of strains NC28.1 and NC85.2 were grown in bacterial suspensions until they reached either a density of 1×10^6^ cells per mL (conditions under which bacteria were still plentiful), or for 10 hours longer than that (conditions under which bacteria have just been depleted). In both treatments, cells were then washed from bacterial leftovers and let aggregate on 3% agar gels

### Imaging samples and counting cells

An ultrasonic atomizer (from CVS) was used to uniformly apply a 0.5mm thick layer of warm imaging solution (SorMC containing 0.3 mol/L of dextrose and 0.05% agar) to the samples. After resting for 10 minutes at room temperature cells assume a spherical shape and detach from neighboring cells. Samples were then photographed with a Canon t5i DSLR camera at a Nikon Ti-Eclipse inverted microscope equipped with a 10X objective. The imaged area, a square of side 1.5cm, was large enough to encompass the initial cell plated area plus a buffer zone that ensures that all cells were imaged. Custom software using the OpenCV package for Python was then used to count the cells that did not join aggregation centers.

The error of automatic cell counts was estimated by taking samples of images of cells in various densities, manually and automatically counting them and fitting a linear Gaussian model with slope 1 and intercept 0 (Figure S2b). The standard deviation of the fitted model was taken as the automatic counting error.

Total plated cell counts in each sample were estimated by counting the non-aggregated 5% dilution of the corresponding stock suspension and multiplying by the dilution factor of that particular sample. We do not assess differential loner viability, which might effectively increase the difference in loner allocation between strains.

### Spatial pattern analysis

For each cell in each experiment, the local cell density was calculated in a neighborhood with a radius of 10% of the total experimental radius. Cells that weren’t within the center of the experimental area (defined by a radius of 60% of the total experimental radius) weren’t included in the analysis, to avoid border effects. For each strain and agar concentration, neighborhood densities of cells of all experiments were pooled together in a probability distribution. The results were qualitatively similar for other neighborhood radii, but because the imaging treatment might shift a bit the position of the cells, we chose a radius that is considerably larger than this effect

### Statistical analyses of mixed strain experiments

Given a constant initial cell density, we intend to test if the loner-aggregator partitioning process of a strain is influenced by the genotypic identity of its neighbors. We let *P*_*i*_ be the proportion of cells of strain *i* that stay as loners. We determined that this depends on *ρ*_0_, the initial density of the population, such that the density of loners in a clonal population of strain *i* at density *ρ*_0_ is 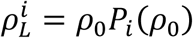. Here we investigate whether in mixes *P*_*i*_ is also a function of the fraction *Π*_*j*_ = 1 *– Π_i_* of cells of co-occurring strain *j* (where *Π_i_* is the fraction of cells of strain *i*). The total loner density in the mix can be expressed as 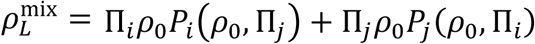, where *ρ*_0_ is the initial total density of plated cells and *Π_i_* is the fraction of strain *i* in the initial mix.

#### Null hypothesis

If *P*_*i*_ is not a function of *Π*_j_, then 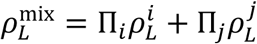, which is the linear combination of the expected loner counts for each of the strains composing the mix (null hypothesis).

#### Statistical test 1

*Piecewise linear regression* (Fig. 3). We measure departures from this linear expectation by fitting a piecewise linear regression to the data. The p-value shows how often data drawn from the global linear fit generates piecewise linear regressions with more extreme inclinations.

#### Statistical test 2

*Maximum likelihood* (Fig. S6a-c). We also use a maximum likelihood based model selection to test for non-linearity. We let 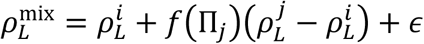 where *f* is the function of interest and *ε* is a normally distributed noise term. If *f*(*Π_j_*) = *Π*_*j*_ then we recover the null hypothesis. In addition, for *f* we explore three other functional forms: sigmoidal, convex and concave, which are given by a shape parameter a as follows:

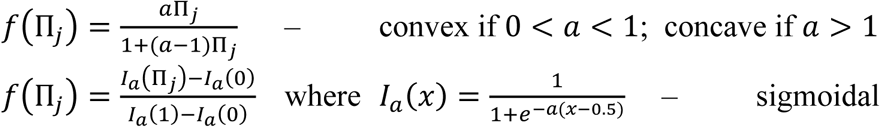

We also considered three forms for the noise term: a homoscedastic structure, a constant coefficient of variation and a heteroscedastic structure, given respectively by

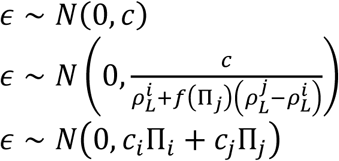

We computed the *Δ*AIC, the difference in AIC (Akaike Information Criterion) between a given model and the best model in the candidate set. Credible intervals were built for the shape parameter *a* using Log-likelihood ratios.

#### Statistical test 3

*Bootstrapping analysis* (Fig. S6d,e). For each of the five strain mix proportions, empirical distributions were bootstrapped and 50.000 data sets were constructed. For each resampled data set, a linear regression was performed using only the pure strain experiments and another linear regression was performed using only the mixed strain experiments. The difference between these inclinations is a measure of the non-linearity of the data set.

### Theory. Population-partitioning model

We implement the spatially-explicit developmental model on a square system of lateral length *ℓ* = 0.2cm that represents a single aggregation territory. Time is discretized in short intervals of length *dt* = 0.01h; our results are independent of the value of *dt*. Within each time step, the internal state and the position of every cell can be updated. Since reproduction and death are negligible over the temporal scales of aggregation, the total population size is conserved during each run of the model.

Consistent with the experimental setup, we initialize the simulations immediately after resource exhaustion with *N*_*0*_ discrete and randomly distributed cells, assumed to be in a pre-aggregating state, *P.* Thus the initial density of cells is *ρ_0_* = *N*_*0*_/*ℓ*^2^. *P-*cells do not move; they emit signal at a constant strain-specific rate *γ* and sense it with a strain-specific sensitivity threshold θ. The assumption that *P*-cells do not move is consistent with experimental results showing a reduced movement of vegetative cells at high density (D’Alessandro et al., 2018). Within each time step *dt*, *P*-cells that sense a local signal density higher than the strain-specific sensitivity threshold *θ,* may become aggregating *A*-cells with a strain-specific probability *λdt*. A detailed description of how signal density is obtained at the position of each cell is below.

*A-*cells move in the direction of the aggregation center, which is exogenously imposed in the center of the system, making a straight displacement of length *vdt* in every time step. This movement pattern simplifies the complexities of *D. discoideum* motion during aggregation, such as the tortuosity in single-cell trajectories caused by imperfect chemotaxis (Fisher, Merkl, & Gerisch, 1989). However, the net effect of these ingredients is to increase the time required to reach the aggregation center, which can be incorporated in the model by changing the value of cell velocity. *A-*cells stop sensing signal but they continue to emit it at the same strain-specific rate, *γ*. When *A-* cells cross the location of the aggregation center (center of the system) in one of their displacements, they adhere to the mound and become multicellular, *M-*cells. *M-*cells do not move and they neither emit nor sense signal. Both the *A*-to-*P* and the *P*-to-*M* transition between cell states are irreversible.

Simulations are allowed to run until the time between two consecutive cell arrivals to the mound is larger than a fixed value *t*_*arr*_ = 1 hour. Alternatively, we explore the effect of a fixed aggregation time by finalizing the simulations after an exogenously imposed time (results not shown). Neither qualitative nor quantitative differences were observed between these two ending conditions, provided that both of these times were sufficiently large. We compute the final density of loners by counting the number of cells that do not belong to the mound at the end of each model realization and dividing it by the area of the system, *ℓ*^2^. Due to the low skewness of the distribution of loner densities obtained from independent realizations, we use the mean loner density, which is obtained by averaging over 100 realizations. A summary of the model parameterization is provided in Table S1.

#### Computation of signal density

Signal is released by both *A*- and *P*-cells, but it is sensed only by *P*-cells. The signal density, *σ*, at time *t* at the position ***r*** of a focal *P*-cell is

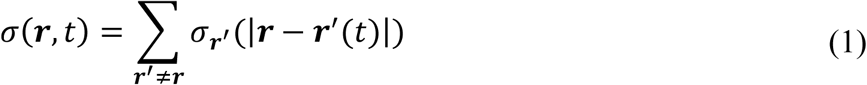

where the index of the sum runs over the locations of all other *A*- and *P*-cells in the system, *|****r*** *–* ***r***^***′***^(*t*)*|* is the distance between the focal cell and these other cells, and 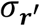 gives the individual contribution of a cell at location ***r***^***′***^ to the total signal density. Since *P*-cells do not move, ***r*** does not depend on *t*; similarly, ***r***^′^(*t*) is either constant (if it is the position of another *P*-cell) or not (if it is the position of an *A*-cell). Our assumptions that signal is continuously released by each cell at a strain-specific rate *γ,* diffuses in the system with diffusion constant *D* and spontaneously decays at rate *η*, lead to a stationary profile in which signal density decreases with the distance from the emitter (see SI Appendix for a detailed calculation of this profile

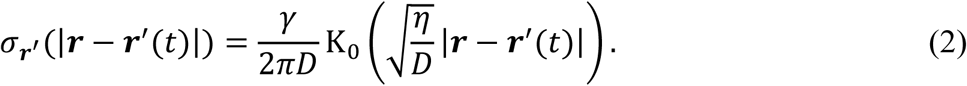

*K*_*0*_ is the zero-order modified Bessel function of the second kind. Since in the experiments aggregation occurs simultaneously on several adjacent aggregation territories, signals may diffuse from one territory to another. To allow for this possibility in the model, the distances between the sensing focal *P-*cell and each of the emitters are measured using periodic boundary conditions.

### Competition model

This model consists of a sequence of growth-starvation periods with the population partitioning between loners and aggregators occurring at the onset of starvation. The expected length of the starvation periods (i.e., mean starvation time) defines the environmental conditions. We discretize each environment into *#* =10^4^ isolated patches (no cell dispersal between them) of area 1 and identical environmental conditions. The model architecture broadly follows (Martínez-García & Tarnita, 2016) but the population partitioning between loners and aggregators is modified to incorporate the behavior produced by the developmental model described above. We assume that the loner-aggregator partitioning curve is constant for all starvation events. Therefore, we do not consider mutation or horizontal gene transfer, which could alter strains’ aggregation behavior.

#### Growth

During growth, free-living amoebae of two different strains compete for a shared resource within each patch. The initial frequency of each strain in the mix is drawn from a standard log-normal distribution, and the total initial population size *X*_*0*_ is normalized to the size of the resource pulse, *R*_*0*_. The size of the resource pulse is fixed and large to guarantee that the population of cells in the patch is also large and that its fluctuations do not affect the aggregator-loner partitioning (i.e., aggregation occurs for population sizes that lie in the region in which loner density plateaus in Fig. S4). Mathematically, the growth dynamics is given by a Monod-like equation:

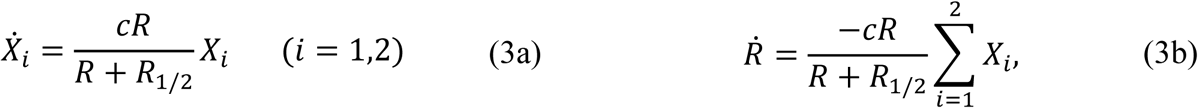

where the dot indicates the time derivative, *X*_*i*_ is the population size for strain *i*, *R* is the amount of resources, and *R1/2* is the abundance of resources at which the growth rate is half of its maximum Here, for simplicity, we assumed that both strains have the same maximum growth rate, although relaxing this assumption constitutes an important expansion towards a more complete understanding of life history traits and tradeoffs in slime molds (Martínez-García & Tarnita, 2016, 2017; J.B.B. Wolf et al., 2015). The growth phase finishes when resources are below a starvation threshold (*R*^*^=1) that is exogenously imposed, since *R* only tends to zero asymptotically in Eq. (3b).

#### Population partitioning

Since cell death is negligible over the temporal scales of aggregation, we assumed that the aggregator-loner partitioning occurs instantaneously upon resource exhaustion. We explored two different scenarios:

(a) Well-mixed development. Each patch is occupied by a homogenous mix of the two strains (Figure S8). Upon resource exhaustion, the density of loners left behind by each of the two strains is determined from the pair-specific co-developmental curve obtained via simulations of co-development using the spatially-explicit developmental model above (Fig. S7a,c). Whenever the composition of the mix at the end of a growth period does not coincide with any of the proportions sampled with co-developmental simulations, we estimate the density of loners of each strain with a linear interpolation between the two closest points to the desired proportion. Finally, since both strains are homogeneously distributed across the whole patch of area 1, the number of loners of each strain *i* 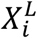 (which is the variable of interest for our model), is identical to loner density. We obtain the number of aggregated cells of each strain as the difference between that strain’s population size upon resource exhaustion (immediately before the population partitioning) and its number of loners.
(b) Segregated development. Each patch is occupied by the two strains but they do not mix; we therefore assume that they occupy a fraction of the patch area equal to that strain’s proportion (Fig. S8). Upon resource exhaustion, the density of loners left behind by each of the two strains is determined from simulations of the developmental model above under clonal conditions. To obtain the number of loners, we multiply the density by the fraction of the patch (of area 1) occupied by that strain. We then obtain the number of aggregated cells as in the well-mixed scenario.

After the population partitioning occurs, based on experimental measures that consistently find an 80:20 spore:stalk ratio within *D. discoideum* fruiting bodies formed under identical conditions (Stenhouse & Williams, 1977), we multiply the number of aggregated cells of each strain *i* by the same constant factor *s* = 0.8 that reflects the effect of spore-stalk cell differentiation. This operation yields the number of reproductive spores, 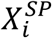.

#### Starvation

The population partitioning is followed by a starvation period of length *T*_*st*_, in which both aggregated and non-aggregated cells die, but at different rates. Spores die at a constant and low rate *δ*, whereas loners have a survival probability, *S*, that decays with time and reaches zero at a maximum survival time *T*_*sur*_. Analogous with (Martínez-García & Tarnita, 2017), we fit this maximum survival time as well as the functional shape of the survivorship curve using experimental data (Dubravcic, 2013) to obtain

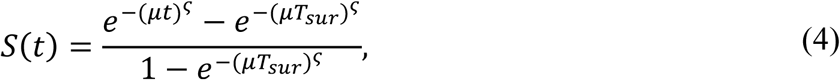

where *μ* is the rate of decrease of the survival probability and *ς* is a parameter that modulates the decay of *S* with time. At the end of the starvation phase, we obtain the populations of surviving loners and spores of each strain *i* as

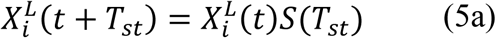

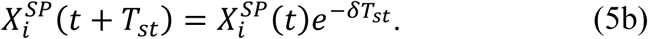

The lengths of the starvation periods *T*_*st*_ are either constant (deterministic environments, in which the length of each starvation period coincides with its mean value) or drawn from an exponential distribution with a mean 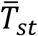 that gives the expected length of the starvation periods. We label deterministic environments using the length of their starvation periods *Tst* and stochastic environments using their expected value 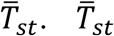.(or *T*_*st*_ in deterministic environments) is a measure for the environmental quality. Lower values of 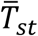 represent better environments in which pulses of resources arrive more frequently on average; larger values of 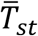 represent worse environments in which resources recover less frequently. Each starvation phase ends with the arrival of the next resource pulse of size *R*_*0*_. We assume that all loners have the same viability regardless of strain (i.e. that different strains do not have differential loner mortality); therefore, upon resource replenishment, all surviving loner cells start reproducing immediately, following Eqs. (3). Spores take an additional time *T*_*ger*_ to germinate (Cotter & Raper, 1968), during which they continue to die at rate *δ*. At the end of the germination time, not all spores become reproducing cells; spores have a probability *ω* of germinating successfully (Dubravcic et al., 2014). Therefore, we multiply the total number of spores by a constant factor *ω* to obtain the fraction of the population of spores that become viable cells and start reproducing according to Eqs. (3).

We repeat this sequence of growth-starvation cycles until one of the strains becomes extinct. We then record the winning strain for each patch and the time-to-extinction for the loser (proxy for transient alpha-diversity). Once we have obtained the winner in each patch, we calculate the variability in the winner across different patches, *Φ* (proxy for stationary beta-diversity), as

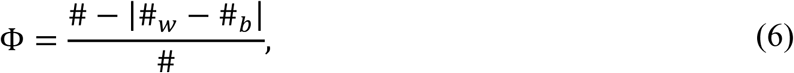

where *#* =10^4^ is the total number of patches, and *#_w_* and *#*_*b*_ are the number of patches dominated by the worse, respectively the better aggregator in the stationary state. From Eq. (6), it follows that *Φ* varies between 0 and 1 (*Φ* = 1 when #*_w_* = #*_b_* and *Φ* = 0 when #*_w_* = # or #*_b_* = #). Since the extinction times and the noise in the extinction times and *Φ* vary depending on the pair of strains and the environmental conditions, mean values are taken over a varying number of independent realizations of the model to optimize computational efficiency. A summary of the model parameterization is provided in Table S1.

## Supplementary Material

**Figure S1.**
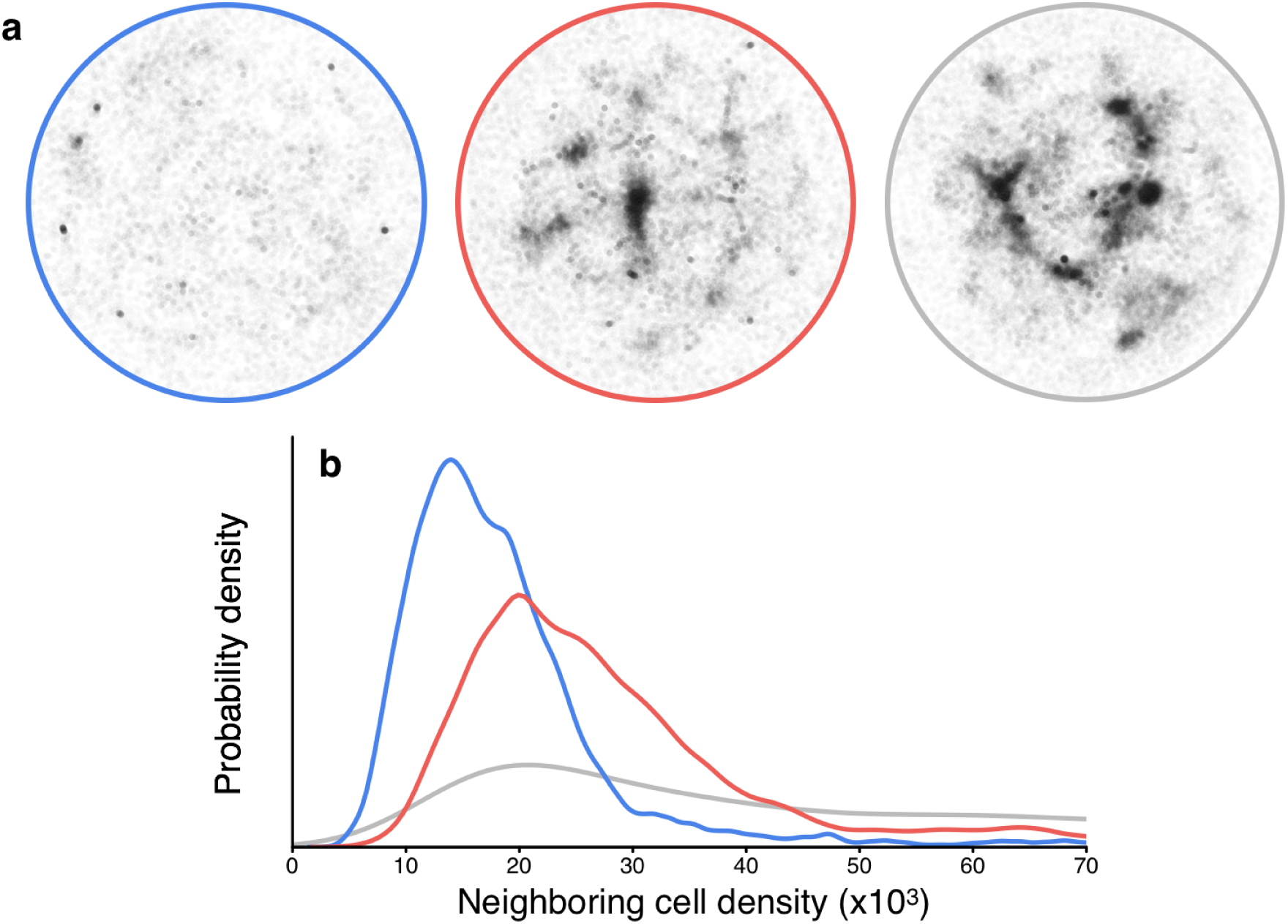
Experimental loner spatial distributions. **a,** Representative loner position maps are shown for each of the three strains (NC28.1 in blue, NC85.2 in red and NC34.1 in grey) plated on 3% agar. The position of each cell is plotted such that darker regions represent regions densely packed with loners. **b,** Characteristic loner spatial patterns for each strain are expressed as the probability distribution of local cell densities (see Methods). Broader peaks and fatter distribution tails (such as for NC34.1) correspond to more heterogeneously distributed loner cells.

**Figure S2.**
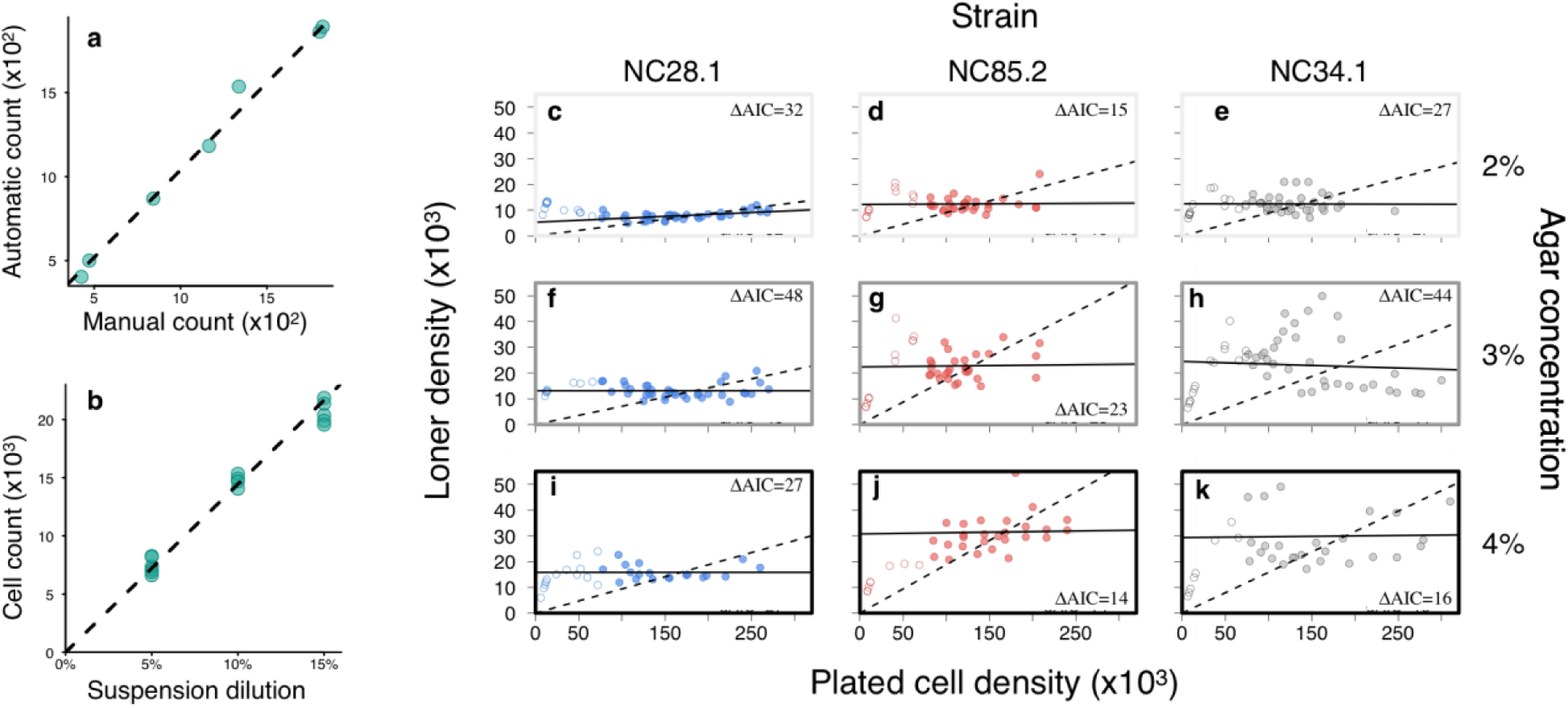
Experimental loner counts. **a,** Loners in regions with varying loner densities were algorithmically counted and plotted against manual (by eye) counts for those same regions. Dashed line = automatic and manual counts coincide. The dispersion around the line is a measure of the counting error. **b,** Cell counts in experiments realized with dilutions from a same cell suspension. Cell densities were below the aggregation threshold. Dashed line = linear regression with intercept anchored at zero. The inclination is a measure of the cell density of the initial suspension, and the dispersion around the regression line is a measure of the error introduced whenever a dilution is made. **c-k,** Loner counts are shown as a function of initial cell plating densities for each of the three strains and each of the three substrate agar concentrations. For initial plating densities above 7.5 x 10^4^ cells/cm2, aggregation occurs for all strains and substrates. To test if above this critical cell density the decision to aggregate is context-independent, those samples with high initial plating densities (filled circles) were used to fit linear Gaussian models with 0 intercept (dashed lines). These zero-intercept models were contrasted to linear Gaussian models with a free intercept parameter (solid lines). *Δ*AIC, the difference in AIC (Akaike Information Criterion) between the 0-intercept and free-intercept models, shows that the latter outperformed the former for all substrates and strains, indicating that the decision to aggregate is context-dependent. Moreover, the inclines of the best fitting linear models are not significantly different from zero for all but the best aggregating conditions (strain NC28.1 on 2% agar substrates), and even then only weakly positive. This indicates that loner densities plateau at high initial plating densities.

**Figure S3.**
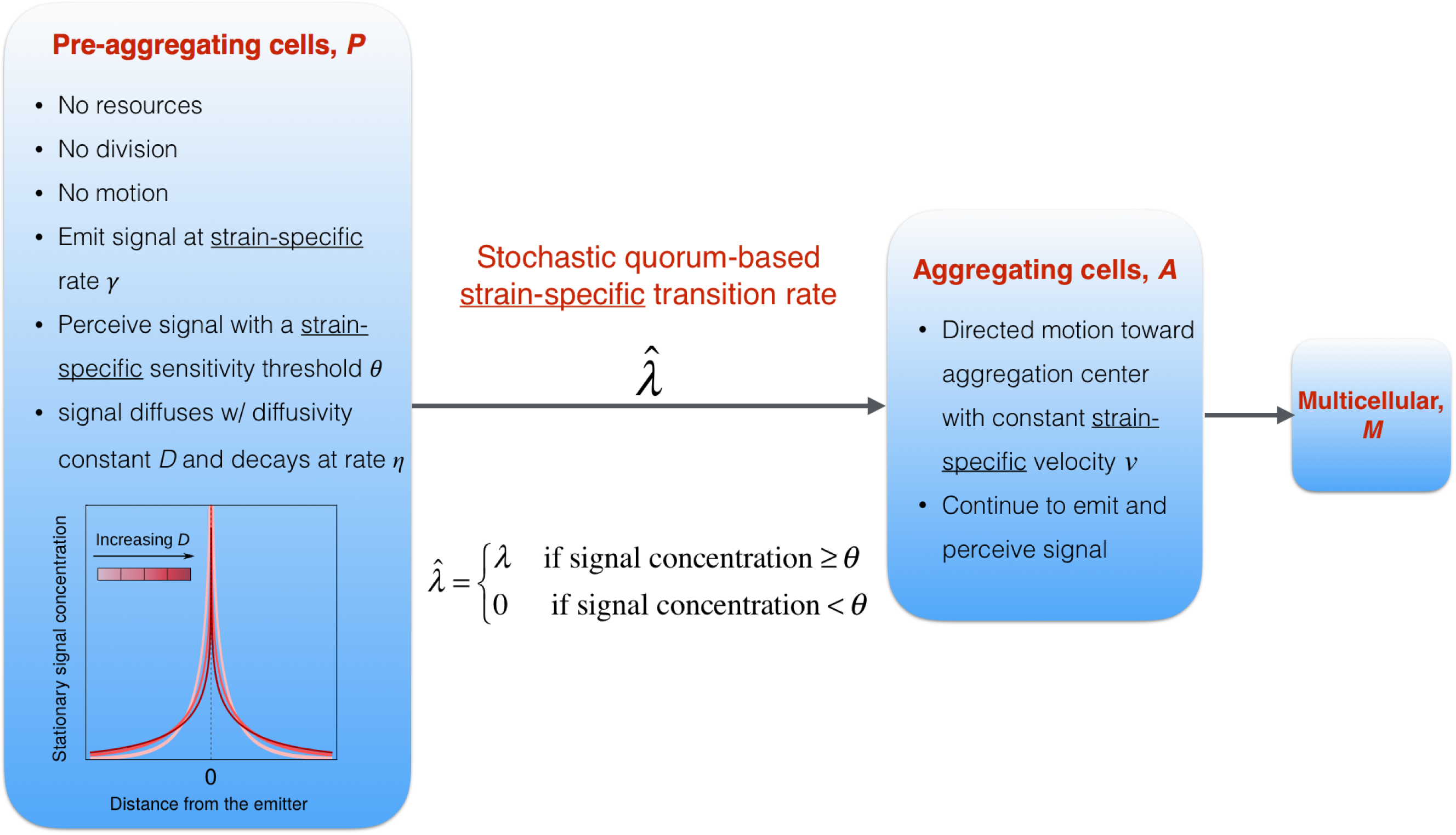
Schematic of the developmental model. We formulated an individual based model approach in which cells can be in three possible internal states: pre-aggregating, *P*; aggregating, *A*; and multicellular, *M*. Each state has different properties, listed in the blue boxes. The transitions between states occur only in one direction, as indicated by the grey arrows. The *P-*to*-A* transition is based on quorum sensing and it occurs at a strain-specific rate, *λ*; for each time step *dt*, if the density of signals is above the strain-specific sensitivity threshold, *P-*cells have a probability *λdt* of becoming *A-*cells. The transition from aggregation to multicellularity is entirely based on movement, and it occurs when cells arrive at the aggregation center.

**Figure S4.**
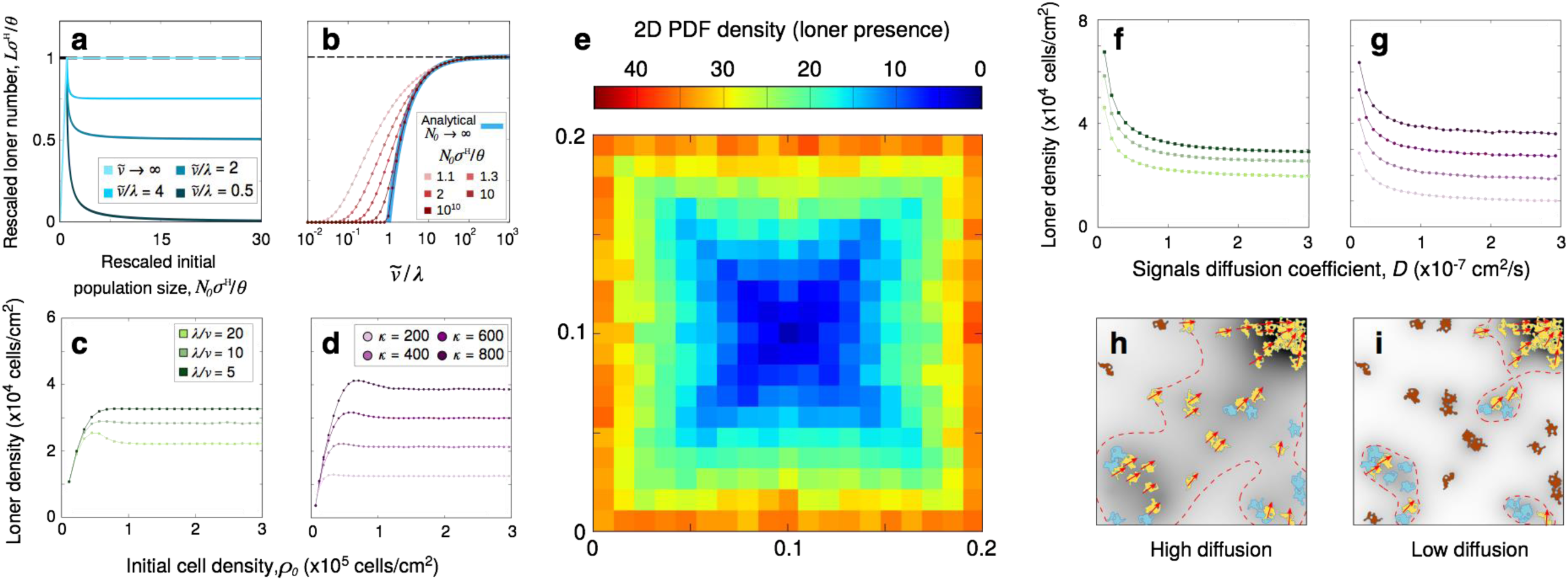
Model results for clonal development. **a**, Analytical solutions in the limit *D* → *∞* for the rescaled number of loners versus the rescaled initial population size (Eqs. (2.13), (2.15), and (2.16) in the SI). **b**, Phase separation between perfectly synchronized (no loners) and asynchronized (loners) development in the limit *D* → *∞*. Red curves are obtained *via* numerical integration of Eqs. (2.11) and (2.12) in the SI for different initial population sizes; the blue-thick curve corresponds to the analytical result in the limit *N*_0_ → ∞ (Eq. (2.22) in the SI and Box 1 in the main text). **c, d**, Loner density versus initial cell density when (**c**) strains differ in λ*/v* with fixed *κ*=500 or (**d**) strains differ in *κ* with fixed λ=1 and *v*=12*μm*/min. *D*=10^-7^. **e,** Probability density function for the presence of loners; the aggregation center is at the center of the system. The histogram is computed using the spatial positions of loners from 100 independent realizations of the model with *D*=3×10^-^8^^, *ρ_0_*=3×10^5^, λ=1, *κ*=400. **f, g,** Loner density versus diffusion coefficient when: **f**, strains differ in λ*/v* with fixed *κ*=500; **g**, strains differ in *κ* with fixed λ=1 and *v*=12*μm*/min. **h, i**, Schematic representation of the reduction in the regions in which signal density is above the strain-specific sensitivity threshold as a result of reducing the diffusion coefficient. Dashed-red lines delineate the regions in which signal density is above a strain-specific sensitivity threshold. Color code for the cells and the concentration of signals as in Figure 2 (a-d). In (**a-g**), non-specified parameters and units are as in Table S1.

**Figure S5.**
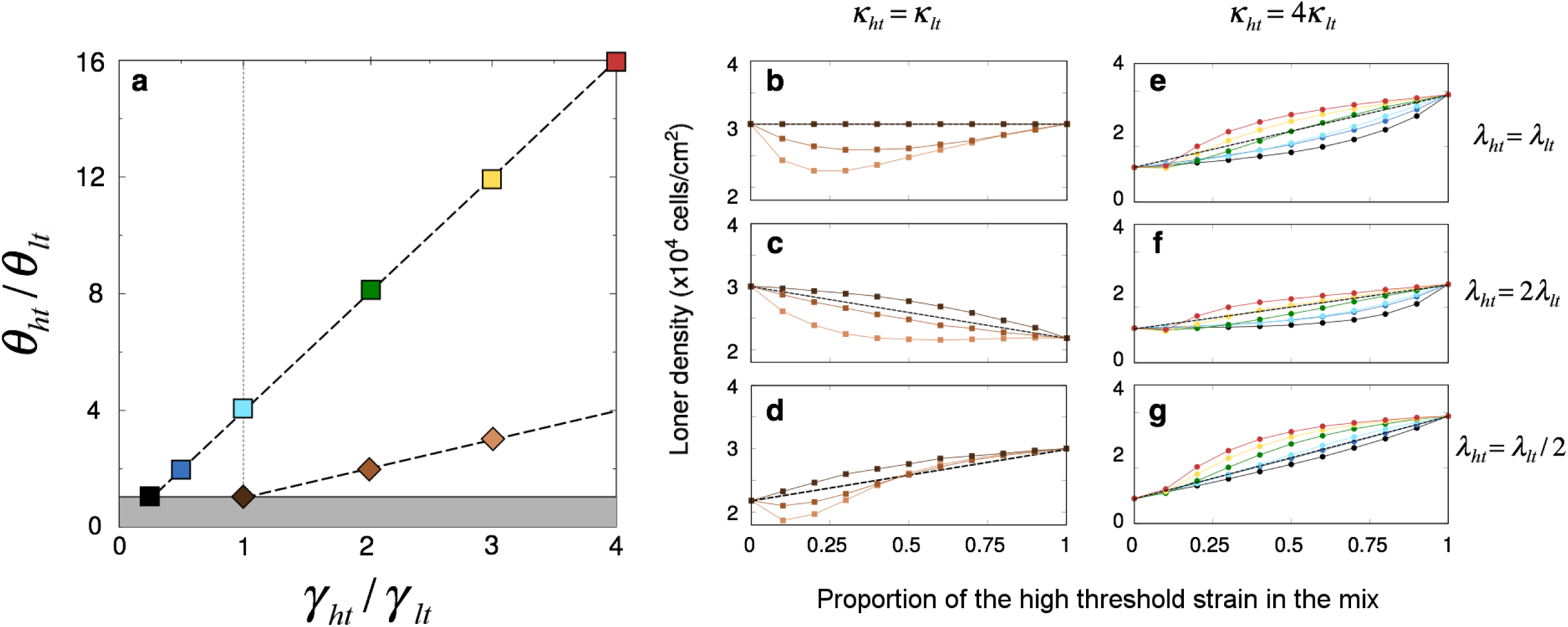
Model results for co-development. For a systematic exploration of the outcome of pairwise developmental interactions within the three-dimensional strain-specific parameter space (*γ, θ, λ*), strains in each mix are labeled according to their relative value of the sensitivity threshold, *θ*. We use the subindex *lt*, standing for ‘low threshold’, to label strain-specific parameter values of the strain with the lowest *θ,* and the subindex *ht*, standing for less sensitive, to label strain-specific parameter values of the strain with the highest *θ*. **a,** γ*_ht_/*γ*_lt_ —* *θ_ht_/θ_lt_* parameter space (*θ_ht_/θ_lt_ >* 1 by definition). The thick-dashed lines trace two transects of the parameter space in which *k*_*ht*_ = *k*_*lt*_ (lower line) and *k*_*ht*_ = 4*k*_*lt*_ (upper line). Densities of mixed loners are shown in (**b-d**) for the parameter values along the lower line and in (**e-g**) for parameter values along the upper line. Specific parameter relationships are indicated by the positions of the squares, whose color is maintained in the mixed-loner curves (**b-g**). **b-d,** *k*_*ht*_ = *k*_*lt*_ = 600, with *θ_ht_* = 300; and *θ_lt_* = 300 (darker brown), *θ_lt_* = 150 (brown), *θ_lt_* = 100 (lighter brown); **b,** λ*_ht_* = λ*_lt_* = 1; **c,** λ*_ht_* = 2, λ*_lt_* = 1; **d,** λ*_ht_* = 1, λ*_lt_* = 2. **e-g,** *k*_*ht*_ = 800, with *θ_ht_* = 400 and *κ*_*lt*_ = 200 with *θ_lt_* = 25, 33, 50, 100, 200, 400 from top to bottom curve (red to black); **e,** λ*_ht_* = λ*_lt_* = 1; **f,** λ*_ht_* = 2, λ*_lt_* = 1; **g,** λ*_ht_* = 1, λ*_lt_* = 2. Dashed lines in (**b-g**) indicate the null hypothesis. Model parameterization shown in Table S1 with *D =* 10^-^7^^ and *ρ_0_* = 3×10^5^. Averages taken over 100 independent model realizations.

**Figure S6.**
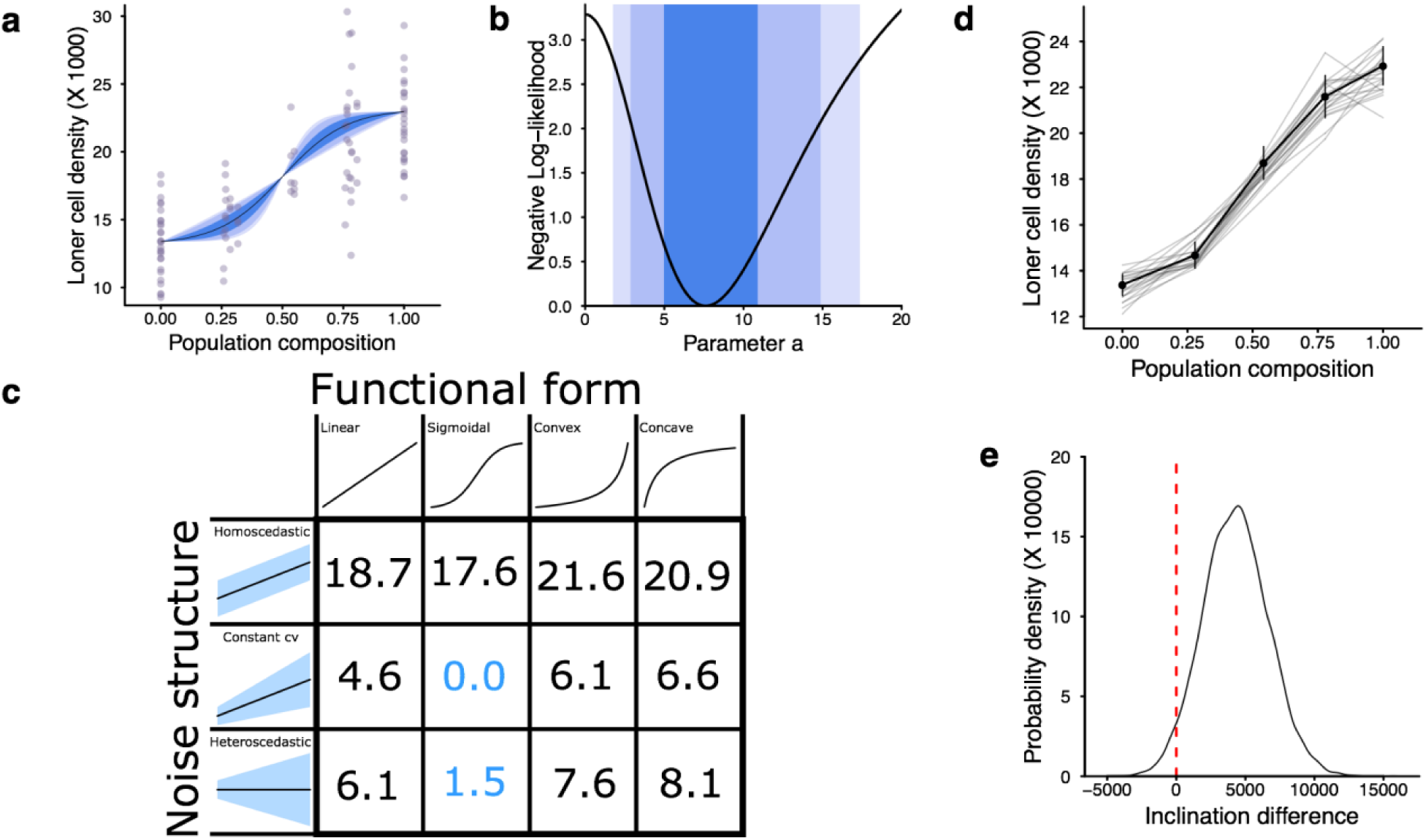
Statistical analysis of non-linearity in mixed strain experiments. a-c, Maximum likelihood analysis. **a,** Grey points = experimental mixed loner densities (see Fig. 3b). Black curve = expected loner densities for the maximum likelihood estimate of shape parameter a (see Methods). Blue areas = envelopes for the loner density curves for the confidence intervals defined by likelihood ratios of 2, 8 and 16, from darker to lighter. **b,** Negative Log-likelihood profile for the shape parameter a of the model with the best Akaike Information Criterion (AIC). Blue areas = confidence intervals defined as in (**a**). **c,** *Δ*AIC, the difference in AIC between a given model and the best model in the candidate set. Blue values = the two best-fitting models. **d,e, Bootstrapping analysis.** For each of the five strain mix proportions, empirical distributions were bootstrapped and 50.000 data sets were constructed. **d,** Grey lines = piecewise linear regressions of 20 of these resampled data sets. Black line = the mean of all resampled data sets. Error bars = standard errors. **e,** For each resampled data set, a linear regression was performed using only the pure strain experiments and another linear regression was performed using only the mixed strain experiments. The difference between these inclinations is a measure of the non-linearity of the data set. Black line shows the probability density function of these inclination differences. Red line at zero marks linearity (p=0.033).

**Figure S7.**
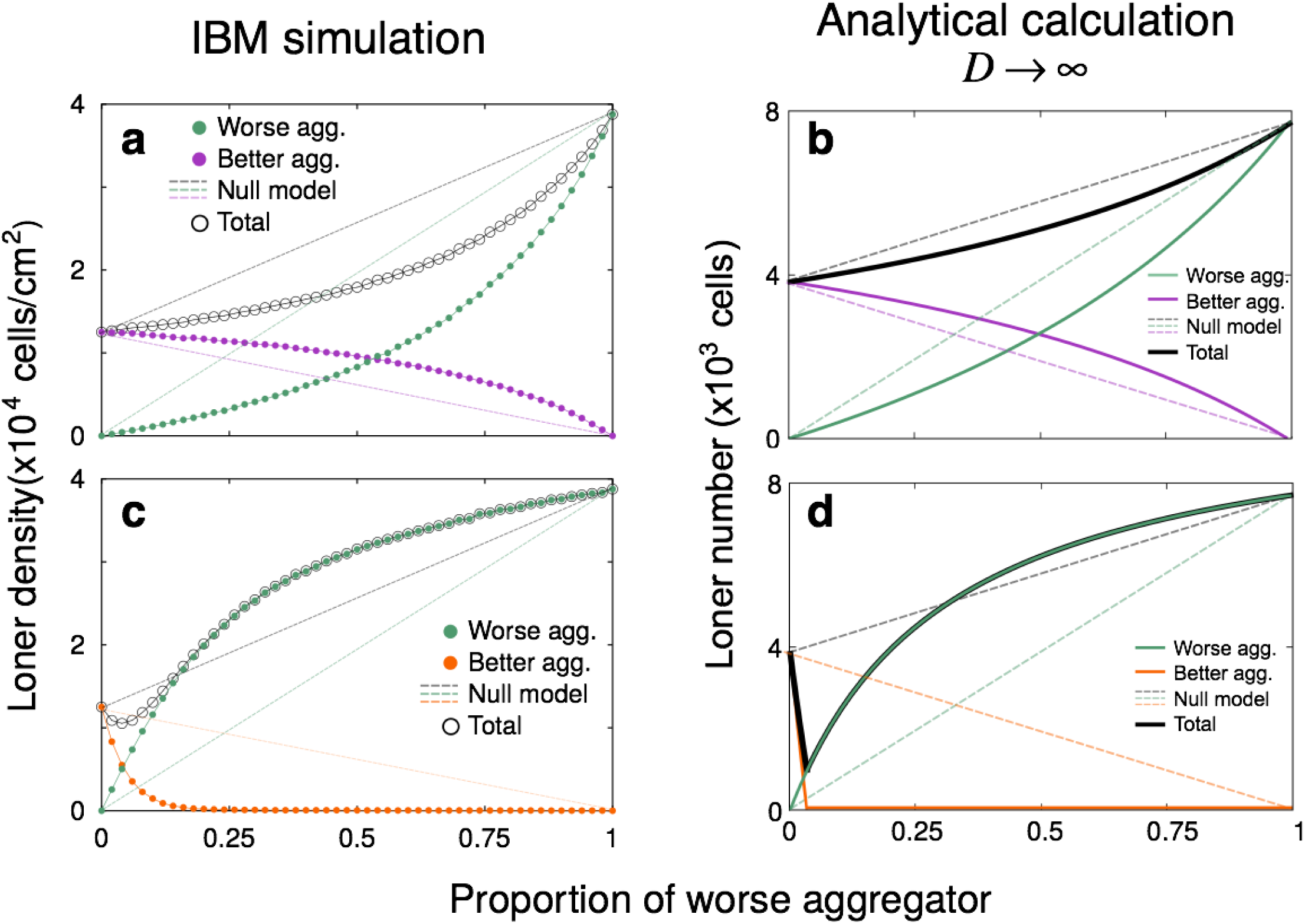
Model results for the effects of co-development on individual strains. Developmental interactions lead strains to become more similar (**a, b**) or more different (**c, d**). (**a, c**) Simulations of the individual based model, *D* =10^-7^. (**b, d**) The analytical approximations to (**a, c**) obtained in the limit *D* → *∞* (Eqs. (2.31) and (2.33) in the Supplementary Information), qualitatively recapitulate the behavior of mixed-loners and of the loners of each strain. Parameterization: *γ_w_* = 0.5, *θ_w_* = 400 (*κ_w_* = 800), λ*_w_* = λ*_b_* = 1, *κ_b_* = 200 with (**a**) *γ_b_* = 1 and (**b**) *γ_b_* = 0.25. *w* = worse aggregator; *b* = better aggregator. Remaining parameters are as in Table S1. The color code for each strain corresponds to Fig. 4.

**Figure S8.**
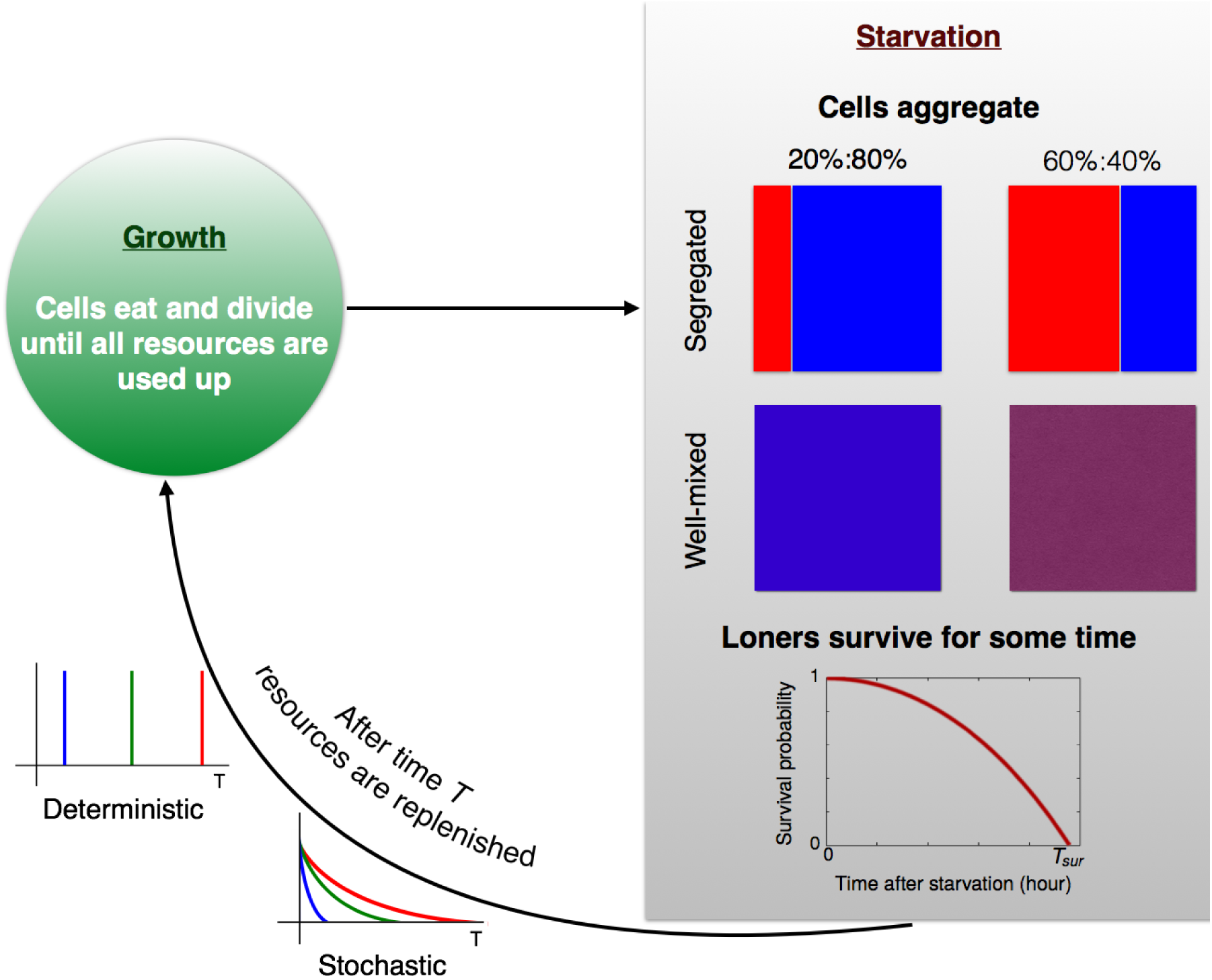
Schematic of the competition model. The model consists of a sequence of growth-starvation cycles. During growth, cells consume a shared pulse of resources and divide; during starvation, loners and aggregated cells die at different rates. The length of the starvation periods *T*_*st*_ can be either fixed (deterministic environments, defined by *T*_*st*_) or drawn from an exponential distribution (stochastic environments, defined by the mean starvation time 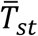). Upon resource exhaustion (at the end of the growth period), the population partitions into aggregators and loners according to our population partitioning model. We compare two scenarios: *well-mixed*, where co-occurring strains co-develop and loner densities are obtained from co-development curves (e.g., as in Figure S7), or *segregated*, where strains are assumed to not mix and loners are derived from each strain’s clonal development partitioning.

**Table S1.**
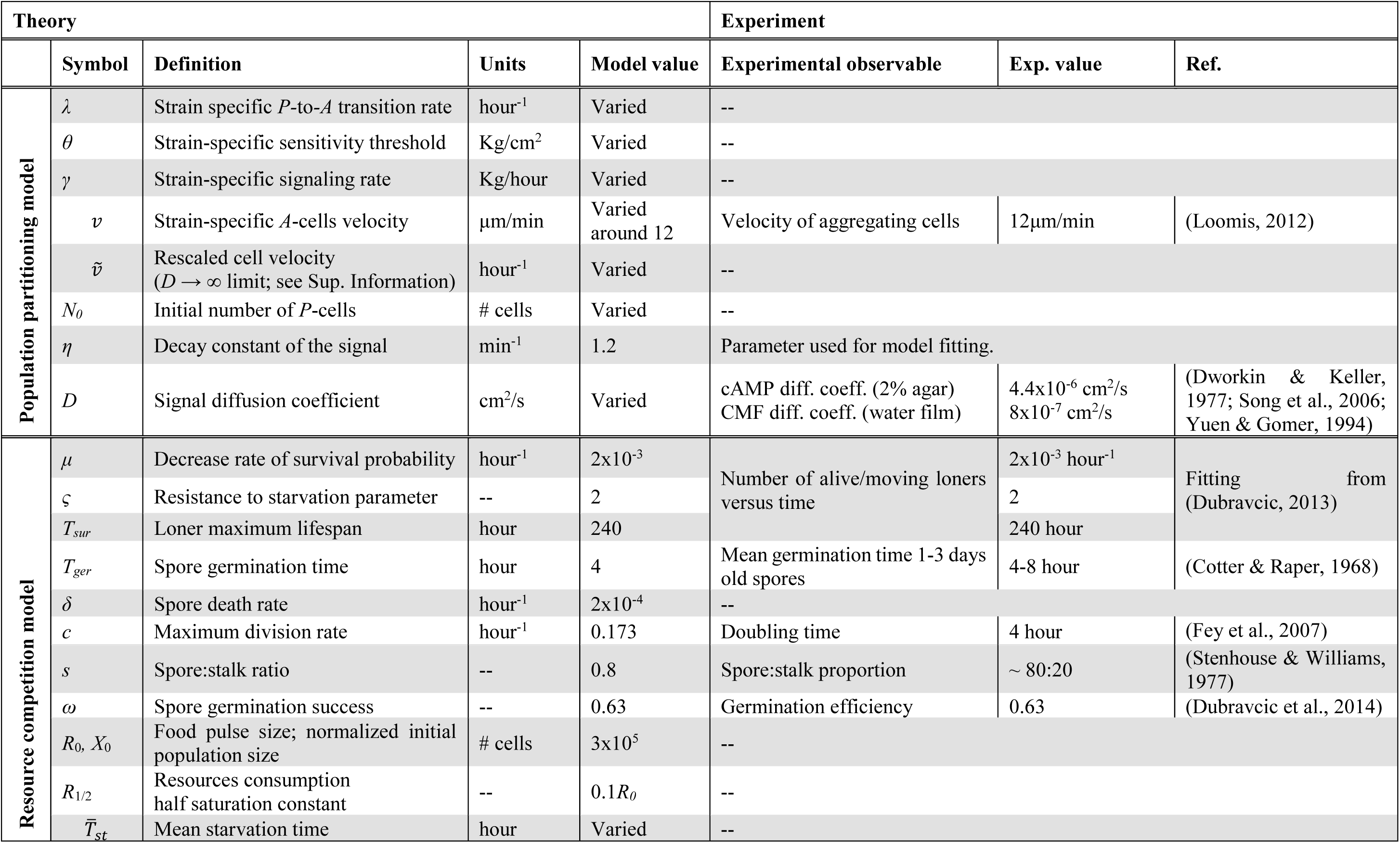
Parameterization of the two theoretical models. CMF is conditioned medium factor.

| Theory |  |  |  |  | Experiment |  |  |
| --- | --- | --- | --- | --- | --- | --- | --- |
|  | Symbol | Definition | Units | Model value | Experimental observable | Exp. value | Ref. |
| Population partitioning model | $\lambda$ | Strain specific *P*-to-*A* transition rate | hour<sup>-1</sup> | Varied | -- | | |
| | $\theta$ | Strain-specific sensitivity threshold | Kg/cm<sup>2</sup> | Varied | -- | | |
| | $\gamma$ | Strain-specific signaling rate | Kg/hour | Varied | -- | | |
| | $v$ | Strain-specific *A*-cells velocity | μm/min | Varied around 12 | Velocity of aggregating cells | 12μm/min | (Loomis, 2012) |
| | $\tilde{v}$ | Rescaled cell velocity ($D \rightarrow \infty$ limit; see Sup. Information) | hour<sup>-1</sup> | Varied | -- | | |
| | $N_0$ | Initial number of *P*-cells | # cells | Varied | -- | | |
| | $\eta$ | Decay constant of the signal | min<sup>-1</sup> | 1.2 | Parameter used for model fitting. | | |
| | $D$ | Signal diffusion coefficient | cm<sup>2</sup>/s | Varied | cAMP diff. coeff. (2% agar)<br>CMF diff. coeff. (water film) | 4.4x10<sup>-6</sup> cm<sup>2</sup>/s<br>8x10<sup>-7</sup> cm<sup>2</sup>/s | (Dworkin & Keller, 1977; Song et al., 2006; Yuen & Gomer, 1994) |
| Resource competition model | $\mu$ | Decrease rate of survival probability | hour<sup>-1</sup> | 2x10<sup>-3</sup> | Number of alive/moving loners versus time | 2x10<sup>-3</sup> hour<sup>-1</sup> | Fitting from (Dubravcic, 2013) |
| | $\varsigma$ | Resistance to starvation parameter | -- | 2 | | 2 | |
| | $T_{sur}$ | Loner maximum lifespan | hour | 240 | | 240 hour | |
| | $T_{ger}$ | Spore germination time | hour | 4 | Mean germination time 1-3 days old spores | 4-8 hour | (Cotter & Raper, 1968) |
| | $\delta$ | Spore death rate | hour<sup>-1</sup> | 2x10<sup>-4</sup> | -- | | |
| | $c$ | Maximum division rate | hour<sup>-1</sup> | 0.173 | Doubling time | 4 hour | (Fey et al., 2007) |
| | $s$ | Spore:stalk ratio | -- | 0.8 | Spore:stalk proportion | ~ 80:20 | (Stenhouse & Williams, 1977) |
| | $\omega$ | Spore germination success | -- | 0.63 | Germination efficiency | 0.63 | (Dubravcic et al., 2014) |
| | $R_0, X_0$ | Food pulse size; normalized initial population size | # cells | 3x10<sup>5</sup> | -- | | |
| | $R_{1/2}$ | Resources consumption half saturation constant | -- | 0.1$R_0$ | -- | | |
| | $\bar{T}_{st}$ | Mean starvation time | hour | Varied | -- | | |

## 1 Calculation of stationary signal profile

We assume that cells are punctual sources that release signal at a constant strain-specific rate *γ*. The signal has a spontaneous decay rate *η* and a diffusion coefficient *D*. Given these conditions, the equation that governs the spatiotemporal evolution of signal density, *σ*(*x, y*; *t*), is

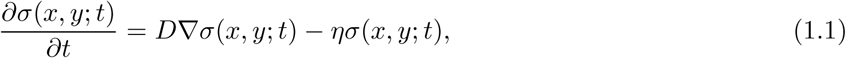

The first term on the right side of Eq. (1.1) accounts for the diffusion of signal and the second term for its spontaneous decay. Here, we first solve the stationary limit (*∂_t_* = 0) of Eq. (1.1) in an infinite domain, imposing as boundary conditions the facts that cells continuously release signals and that signal density goes to zero when the distance from the emitting cell tends to infinity. Subsequently, we discuss the effect of considering a finite integration domain with periodic boundary conditions.

Due to the radial symmetry of the problem, we transform Eq. (1.1) to polar coordinates, in which the partial differential equation in (*x, y*) becomes an ordinary differential equation in the radial coordinate *r* that indicates the distance to the source of the signal,

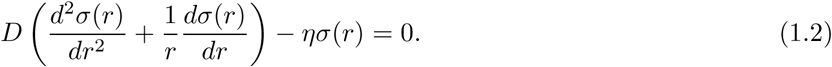

Since the position of the emitter, *r* = 0, is a singular point of Eq. (1.2), we will first assume that cells have a finite radius 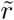 and then take the limit 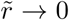 After the transformation to polar coordinates, and assuming a finite radius for the cell, the boundary conditions can be written as,

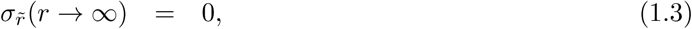

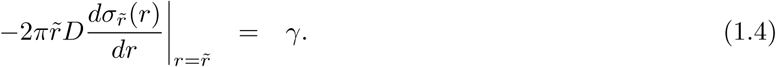

Eq. (1.3) imposes the finiteness of the density, and Eq. (1.4) imposes that the amount of mass released through the boundary of the cell per unit time has to be constant and equal to the strain-specific emission rate *γ*. The subscript 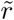 in the densities accounts for the finite radius of the cells.

Equation (1.2) is the modified Bessel equation of order zero, and its general solution can be expressed as

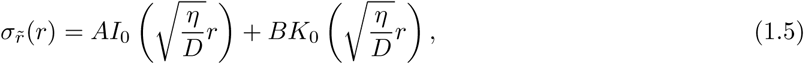

where *I*_0_ and *K*_0_ are the zero order modified Bessel functions of the first, respectively second, kind. From the boundary condition of Eq. (1.3), it follows that *A* = 0, since *I*_0_ diverges when its argument tends to infinity. *B* is calculated from the second boundary condition, Eq. (1.4),

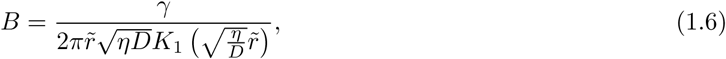

where *K*_1_ is the first order modified Bessel function of the second kind and we have used that 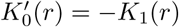.

Inserting Eq. (1.6) into (1.5), the stationary signal profile produced by a source of finite radius 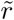 is,

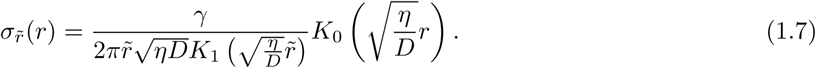

Finally, to obtain the profile generated by a punctual source, we take the limit 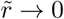 in Eq. (1.7),

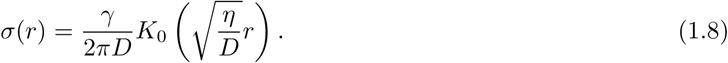

The solution provided by Eq. (1.8) assumes an infinite system size, whereas we perform numerical simulations of the developmental model on a finite domain of lateral length *ℓ* with periodic boundary conditions. To impose periodic boundary conditions is equivalent to considering that the simulated finite domain corresponds to a tile embedded into an infinite lattice in which each tile is a mirroring image of the focal domain. The signal density within the focal tile is obtained by adding over the contributions of all other tiles. However, since our numerical simulations only explore a range of diffusion coefficients in which *σ*(*l/*2) ≈ 0, we can truncate the sum over tiles at the nearest neighbors of the focal one. This is equivalent to calculating distances to the position of each emitting cell, (*x*_em_, *y*_em_), in each spatial coordinate:

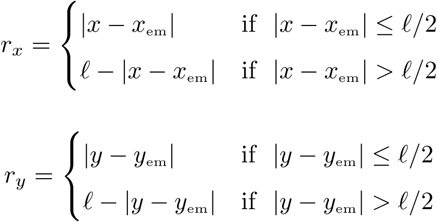

The total distance is then given by the radial coordinate *r*, as 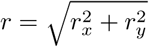.

## 2 Analytical treatment of the developmental model in the spatially-implicit limit *D* → ∞

The spatially-implicit limit of the individual based population-partitioning model consists of disregarding the spatial effects introduced by a finite signal diffusion coefficient (i.e. the limit *D* → ∞), but still accounting for cell movement at a finite velocity. To this end, we map cell movement into a stochastic transition in cell state from aggregating to being multicellular; the rate of this transition is related to cell velocity, *v*.

First, we calculate the stationary signal density profile produced by each cell in the limit *D* → ∞. Unlike in the low *D* case explored in the spatially-explicit simulations, in which periodic boundary conditions were implemented considering only the nearest neighbors of the focal tile, now, since the signal spreads infinitely far, we need to include the contribution of an infinite number of tiles. This results in each cell generating a homogeneous signal distribution within the focal tile, *σ*^H^ = ℳ*/l*^2^. ℳ is the mass of signal that is being released by each cell in the stationary limit, which can be obtained by integrating Eq. (1.8) over the entire range of distances,

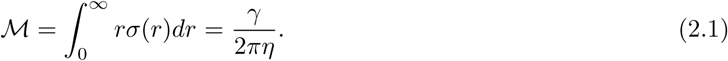

Due to the conservation of the total population size *N*_0_ (since demographic events are neglected on the temporal scales of aggregation), the state of the system is fully determined by the sizes of two of the three subpopulations (*P, A*, and *M* cells). We choose the number of cells in the *P*-state, *N*_*P*_, and in the *A*-state, *N*_*A*_, as state variables. The number of cells in the *M* state, *N*_*M*_ (*t*) (i.e., the size of the multicellular aggregate) can then be obtained from *N*_*P*_ (*t*) + *N*_*A*_(*t*) + *N*_*M*_ (*t*) = *N*_0_.

In order for the aggregation process to be initiated at all, a quorum must be met by the initial population (all of which are *P*-cells), i.e. we must have *N*_0_*σ*^H^ *> θ*. In the absence of a quorum, all initial cells remain as loners and therefore the total loner number is *L* = *N*_0_. If there is an initial quorum, then *P*-cells turn into *A*-cells at rate *λ*; *A*-cells continue to emit signal while they move in the direction of the aggregate. As *A*-cells eventually join the aggregate, they stop signaling and therefore the amount of signal in the system continues to decrease. *P*-cells continue to become *A*-cells at rate *λ* only if the total signal density [*N*_*P*_ (*t*) + *N*_*A*_(*t*)] *σ*^H^ remains above the strain-specific sensitivity threshold, *θ*. The *P*-to-*A* transition rate as a function of time is thus given by

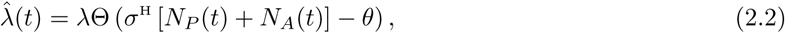

where Θ is the Heaviside function, which takes value 1 for non-negative arguments and 0 for negative arguments. Therefore, omitting the temporal dependence in 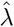, *N*_*A*_ and *N*_*P*_,

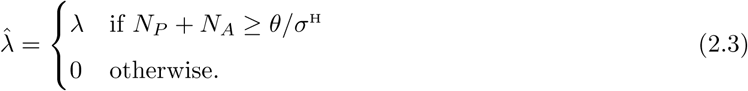

The rate at which *A*-cells stick to the aggregate and become *M*-cells can be approximated by the inverse of the time needed to cover the mean distance to the aggregate at a velocity *v*, i.e.

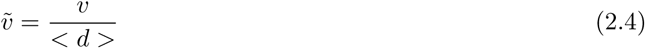

where *< d >* is a characteristic spatial scale of the aggregation territory (mean distance to the aggregation center). For simplicity, we will fix *< d >*= 1 in the following and refer to 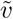 as a rescaled velocity.

Therefore, the aggregation process can be mapped to a sequence of two stochastic reactions, each of which occurs at a different rate,

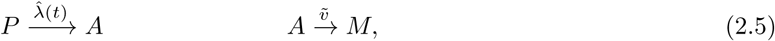

This stochastic process is fully described by a master equation, which gives the temporal evolution of the probability *g*(*N*_*P*_, *N*_*A*_; *t*) of finding the system in a state (*N*_*P*_, *N*_*A*_) at time *t*,

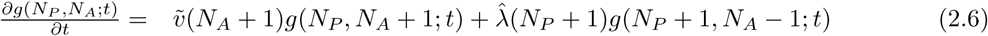

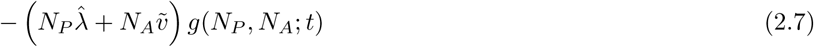

To simplify the notation, the temporal dependence in *N*_*A*_, *N*_*P*_ and 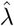 has been omitted.

Following standard procedures, from the Master equation (2.7) we can derive a system of coupled ordinary differential equations for the mean value of each subpopulation size,

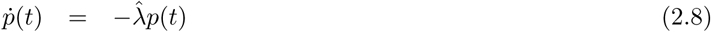

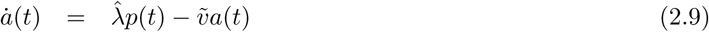

where *p*(*t*) and *a*(*t*) are the mean values of *N*_*P*_, respectively *N*_*A*_, at time *t*. The dot over *a* and *p* on the left side of the equation indicates a time derivative. System (2.8) can be solved analytically, using that initially all cells are in the pre-aggregation state, i.e. *p*(0) = *N*_0_, *a*(0) = 0. Then

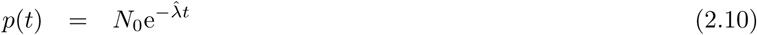

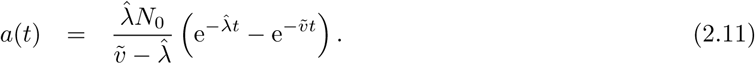

Since the ultimate objective of this approximation is to obtain analytical expressions for the loner-aggregator partitioning behavior, an important observable is the time *τ* at which the decaying signal density exactly equals the strain-specific sensitivity threshold. *τ* can thus be obtained by solving

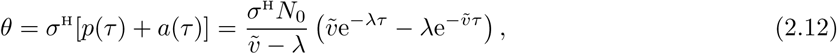

where we have used the fact that 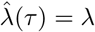 according to Eq. (2.2). Since aggregating cells also contribute to the pool of signal, *τ* does not represent the aggregation time; after a time *τ*, any *A*-cell in the system will continue to move towards the aggregate at rate 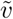 until *a*(*τ* + Δ*t*) = 0. However, importantly, *τ* gives the time at which the last *P* - *A* transition occurs. Therefore, all cells that are still in the *P*-state at time *τ* will remain as loners and we can find the total number of loners as

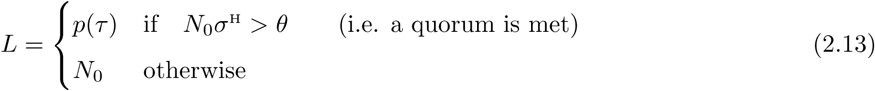

Henceforth we will focus on the former case, when aggregation does get initiated.

In general, we can not solve for *τ* in Eq. (2.12) and therefore we can not determine the number of loners analytically. Below, we try to circumvent this problem by looking at a few special cases.

### 2.1 Analytical results for the non-spatial limit 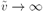

In this limit, cells spend an infinitesimally short time in the *A* state and therefore *p*(*t*) + *a*(*t*) → *N*_0_e^-*λ/t*^. To obtain *τ* we

then solve *σ*^H^ *N*_0_e^-*λ/τ*^ = *θ*, which gives 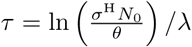. Then, from Eq. (2.13), the number of loners, when there is a quorum for aggregation, is

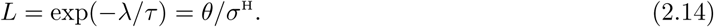

Therefore, in this limit, *λ* gives the time scale of the aggregation but it has no effect on the number of loners, which is equal to the sensing-to-signal ratio.

### 2.2 Analytical results when 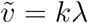 or 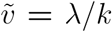

In the special case 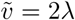, using the change of variables *y* = exp(-*λτ*), (2.12) becomes a quadratic equation from which *y* and thus *τ* can be obtained,

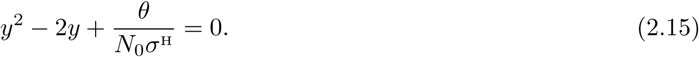

Given the definition of *y*, Eq. (2.15) only has physical meaning in the domain *y* ∈ (0, 1]. Within that interval, if a quorum exists (i.e. *N*_0_ *> θ/σ*^H^), Eq. (2.15) has a single root, which determines the number of loners *p*(*τ*) according to Eq. (2.10) and the definition of *y*:

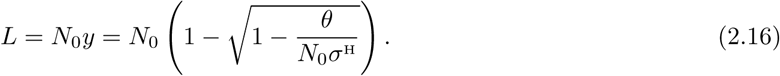

In the limit *N*_0_ → ∞, Eq. (2.16) tends to *θ/*(2*σ*^H^), as predicted by Eq. (1) in the main text (also Eq. (2.23) below).

In the other special case, 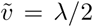, Eq. (2.12) becomes again Eq. (2.15) using the change of variables 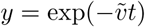. Thus, if there is a quorum for aggregation (i.e. *N*_0_ *> θ/σ*^H^) the number of loners is

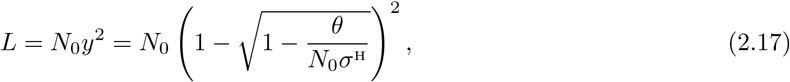

which tends to 0 in the limit *N*_0_ → ∞, as predicted by the phase separation defined by Eq. (1) of the main text (also Eq. (2.23) below).

In general, the changes of variables introduced here, *y* = exp(-*λt*) and 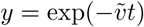 will turn Eq. (2.12) into a polynomial equation of degree *n* provided that 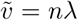 or 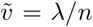. If the root of such a polynomial within the interval *y* ∈ (0, 1] can be obtained, then an expression for the number of loners as a function of the initial population *N*_0_ is accessible. In Figure S4, we show the two cases obtained here (Eqs. (2.16) and (2.17)), as well as the non-spatial limit 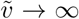 In addition, we also show the 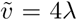 case, where the equivalent to Eq. (2.15) is a 4-th degree polynomial, whose root in the interval *y* ∈ (0, 1] we obtained using *Mathematica 11.1*.

### 2.3 Analytical results in the large population limit *N*_0_ → ∞

In the limit of infinitely large initial population size, a quorum is always met. Therefore, from Eq. (2.13), the number of loners is

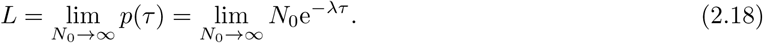

At the end of this section, we prove that *p* is a positive and monotonically decreasing function of *N*_0_; therefore *L* always exists and is greater than or equal to zero. This also implies that 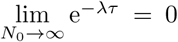 (otherwise *L* would not be finite); applying L’Hôpital’s rule to Eq. (2.18), we obtain

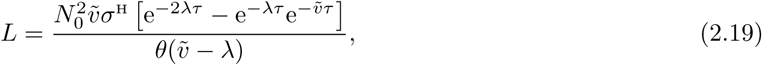

where we have used Eq. (2.26) for the derivative of *τ* with respect to *N*_0_. Finally, defining 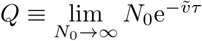 and rearranging terms, we get

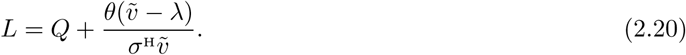

To obtain an independent expression for *L* we need another, non-redundant relationship between *L* and *Q*. This can be obtained by first rearranging terms in Eq. (2.12),

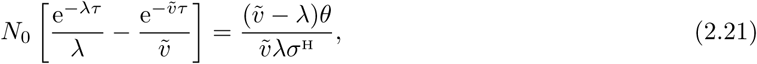

and then taking the limit *N*_0_ → ∞ in Eq. (2.21),

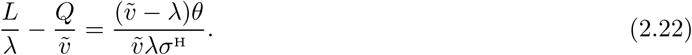

Solving for *L* in Eqs. (2.19) and (2.22), we find

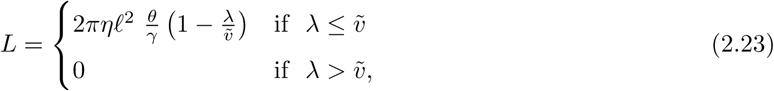

where we have used the fact that 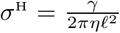 (see Section 2). In the limit *N*_0_ → ∞, there is thus a phase separation given by the relative magnitudes of the *P*-to-*A* and *A*-to-*M* transition rates.

***Proof of the existence of L***. In order to obtain in Eq. 2.23 the limit of *p*(*τ*) for infinite initial population sizes, we first need to prove that such a limit exists and is finite. To this end, we will first calculate the derivative of *p*(*τ*) with respect to *N*_0_:

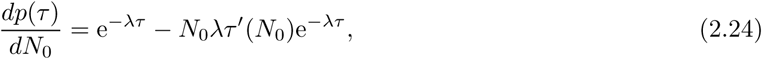

Although Eq. (2.12) for *τ* cannot be solved in general, we can obtain an analytical expression for the derivative of *τ* with respect to *N*_0_ using implicit differentiation. We differentiate both sides of Eq. (2.12)

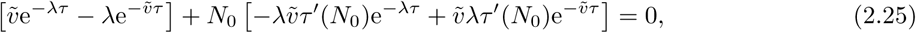

and solving for *τ* ^′^(*N*_0_), we obtain

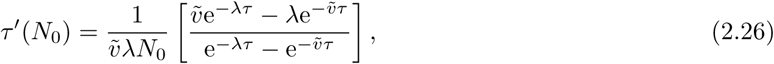

which is always positive for any relationship between 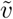 and *λ*. Using Eq. (2.26) in Eq. (2.24), we find

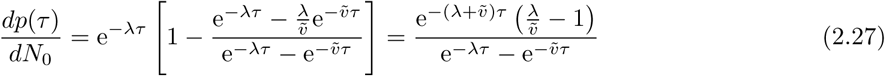

Since the numerator and the denominator of Eq. (2.27) have opposite signs for both 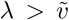 and 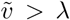, *dp*(*τ*)*/dN*_0_ is always negative. Thus, *p*(*τ*) is a decreasing function of *N*_0_. Since *p* is a non-negative and decreasing function, the limit of *p*(*τ*) as *N*_0_ tends to infinity exists and is always greater than or equal to zero. Importantly, due to the symmetry between *p* and 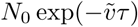, the limit *Q* defined in the calculation of *L* also exists and has the same properties as *L*.

### 2.4 Analytical results for co-development of two strains with same *λ* and 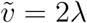

In mixed development, we consider two strains defined by the set of strain-specific parameters (*λ, θ, γ*). *λ* and *θ* have been defined above, and *γ* determines the strain-specific signal density *σ*^H^ released by each cell. We use the term *high-threshold strain* and the notation *ht* for the strain with the higher signal sensitivity threshold and *low-threshold strain* (*lt*) for the one with the lower signal-sensitivity threshold. Thus, *θ_ht_ > θ_lt_*.

If both strains have the same strain-specific *λ*, which is the case studied in this section, then the high-threshold strain also has the higher investment in loners (i.e. it is the worse aggregator) if

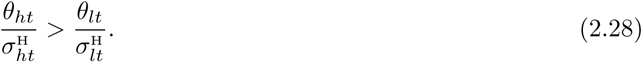

To obtain the mixed loners, we generalize Eq. (2.12) to the two-strain case,

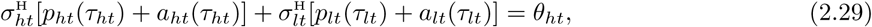

where *τ*_*ht*_ is the time at which the density of signals reaches the strain-specific sensitivity of the high-threshold strain. Therefore, for 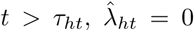 and only cells of the low-threshold strain continue to aggregate 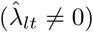.

Let Π be the proportion of the high-threshold strain in the mix; Π*N*_0_ is thus the initial population of the high-threshold strain, and (1 - Π)*N*_0_ the initial population of the low-threshold strain. Substituting the expressions for *a*(*t*) and *p*(*t*) obtained in Eqs. (2.10) and (2.11), Eq. (2.29) becomes

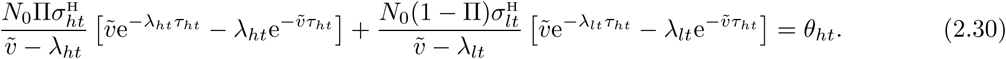

*τ*_*ht*_ cannot be obtained from Eq. (2.30) in general. However, since we assume in this section that *λ*_*ht*_ = *λ*_*lt*_ = *λ* and 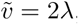, then the change of variables *y* = exp(-*λτ_ht_*) turns Eq. (2.30) into a quadratic equation of the form

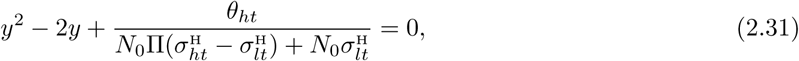

that has only one root in the interval (0, 1]. Using that root, we obtain the number of loners of the high-threshold strain,

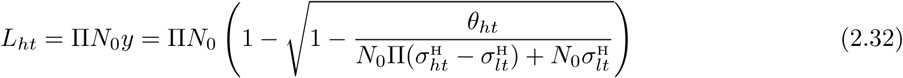

After *τ*_*ht*_, only cells of the low-threshold strain continue to aggregate. The number of loners left by the low-threshold strain is determined by the relationship between 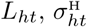, and *θ*_*lt*_:

- If 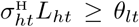 the loners of the high-threshold strain provide quorum for a full aggregation of the low-threshold strain and therefore *L*_*lt*_ = 0.
- If 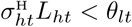, the low-threshold strain stops aggregating at a time *τ*_*lt*_ > τ_*ht*_, such that

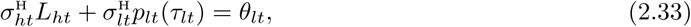

which gives the number of loners for the low-threshold strain

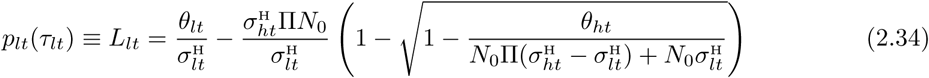 The transition from one outcome to the other occurs at a population composition 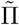 such that 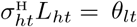 Using Eq. (2.32) for the number of loners of the high-threshold strain, we obtain

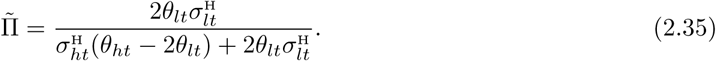

